# Leveraging Bayesian Networks for Consensus Network Construction and Multi-Method Feature Selection to Decode Disease Prediction

**DOI:** 10.1101/2025.04.07.647660

**Authors:** Rosa Aghdam, Shan Shan, Richard Lankau, Claudia Solís-Lemus

## Abstract

Constructing reliable microbiome co-occurrence networks and identifying disease-associated taxa remain major challenges in microbiome research due to variability introduced by different inference algorithms. To overcome these limitations, we present `CMIMN`, a novel R package that uses a Bayesian network framework based on conditional mutual information to infer robust microbial interaction networks. To further enhance reliability, we construct a consensus microbiome network by integrating results from `CMIMN` and three widely used methods—`SPIEC-EASI`, `SPRING`, and `SPARCC`. This consensus approach, which overlays and weights edges shared across methods, reduces inconsistencies and provides a more biologically meaningful view of microbial relationships. In addition, we introduce a multi-method framework for identifying disease-associated microbial taxa by combining machine learning and network-based feature selection. Our ML pipeline applies distinct algorithms and identifies key taxa based on their consistent importance across models. Complementing this, we employ two network-based strategies that prioritize taxa based on centrality differences between ‘clean tubers’ and ‘scab-infected tubers’ networks, as well as a composite scoring system that ranks nodes using integrated network metrics. Our results show that `CMIMN` achieves high robustness in network inference, and that the consensus network further improves stability and interpretability. The multi-method feature selection framework enhances confidence in identifying biologically relevant taxa linked to potato common scab disease. Notably, we identify *Bacteroidota*, *WPS-2*, and *Proteobacteria* at the Phylum level; *Actinobacteria*, *AD3*, *Bacilli*, *Anaerolineae*, and *Ktedonobacteria* at the Class level; and *C0119*, *Defluviicoccales*, *Bacteroidales*, and *Ktedonobacterales* at the Order level as key taxa associated with disease status.

## 1 Introduction

Potatoes, as the world’s fourth most essential crop, play a vital role in addressing global food security. However, soil-borne diseases like potato common scab, caused by *Streptomyces* scabies and related pathogens, significantly threaten potato yield and quality. This disease results in economic losses due to the rejection of tubers with pitted, corky lesions and has broader implications for food security. While some growers and consultants may claim that fumigation with broad-spectrum fungicides controls common scab symptoms, evidence for its effectiveness remains inconsistent, and fumigation is often costly and environmentally restrictive [1]. In practice, genetic resistance (or tolerance) in potato cultivars has emerged as a more effective and sustainable strategy for managing the disease [2]. A promising complementary approach involves exploiting the potential of naturally disease-suppressive soils, which harbor specific microbial communities that suppress pathogens and reduce disease outbreaks. Research has shown that suppressive soils often harbor distinct microbial communities with a higher abundance of antagonistic taxa, such as non-pathogenic *Streptomyces spp.* and *Bacillus* [3, 4]. These insights underscore the importance of investigating microbial interactions in suppressive soils to guide environmentally sustainable disease control practices.

Understanding the composition of microbial communities and the environmental factors shaping this composition is crucial for comprehending biological processes [5–10] and predicting plant phenotypic variations related to plant health and crop production [11–14]. To fully explore the complex interactions between microbes and their environment, we need robust computational approaches that can accurately represent microbial communities. Graphical structures like networks offer a strong mathematical framework for examining organismal relationships across various interactions, including those observed in food webs, plant-plant interactions, plant-animal associations, and their applications in detecting networks within gene regulation and protein-protein interaction systems [15]. Networks provide a formal yet intuitive representation of complex systems, where species are represented as nodes, and their interactions are represented as edges [16]. Although network analysis is widely used in microbiome studies, its application to soil microbial communities has emerged more recently, marked by growing interest in co-occurrence analysis. [17]. However, the complexity of soil presents unique challenges in constructing and interpreting network models, stemming from the need to account for the inter- and intra-variability of samples, which results from the intrinsic heterogeneity of soil conditions [10].

In recent years, the field of microbiology has witnessed significant advancements in network analysis techniques. Studies conducted by Wagg et al. [17] have pioneered the application of network theory to understand complex microbial interactions in various soil microbiome systems. Their findings highlight the potential of community network complexity in influencing ecosystem functions, suggesting that microbial interactions play a crucial role in soil health and resilience. However, their results also indicate that simpler diversity metrics, such as species richness, may explain a substantial proportion of variation in ecosystem functionality. This underscores the need to combine network metrics with traditional diversity measures to obtain a comprehensive understanding of microbial community functions. Moreover, network analysis in microbiome studies requires substantial methodological improvements, as demonstrated by Guseva et al. [15], who showed that different network-construction algorithms can significantly impact the inferred structure of microbial networks in soil ecosystems. This variability highlights the importance of carefully considering methodological choices when applying network analysis to soil microbial datasets, enabling more robust and hypothesis-driven research. In particular, inconsistencies across network inference methods pose a major challenge, limiting the reliability of microbiome-based discoveries.

Working with microbiome data is inherently challenging due to its sparse, high-dimensional, and compositional nature [18]. Numerous methods exist to infer the structure of microbiome networks, but the results often exhibit minimal overlap across different approaches, reflecting the variability and complexity of the data [15]. Furthermore, the absence of a universally accepted gold standard for network evaluation complicates efforts to assess and validate inferred networks. Thus, there is an urgent need for methodological advancements that can enhance the reliability of microbiome network inference and provide biologically meaningful insights.

In addition to constructing reliable networks, identifying important disease-related taxa is a key aspect of microbiome studies, particularly in the context of disease management [19]. Such insights can provide farmers with actionable tools to better understand and control the microbial ecosystems influencing crop health and productivity. Developing robust methods to identify taxa associated with disease resistance or susceptibility is essential for translating microbiome research into practical agricultural solutions. By identifying microbial indicators of plant health, researchers can advance both scientific understanding and practical applications, offering sustainable strategies to mitigate disease impacts and enhance agricultural outcomes.

This study builds upon prior research that explored associations between soil properties and biological phenotypes using machine learning models, including random forests and Bayesian neural networks [18]. Extending this work, we adopt a network-based perspective—specifically leveraging Bayesian networks—to investigate microbial relationships within the soil microbiome. Our approach addresses two primary challenges in microbiome analysis: constructing reliable co-occurrence networks and identifying microbial taxa associated with potato common scab disease. To enhance the robustness of disease-associated OTU identification, we introduce a comprehensive multi-method framework that combines machine learning-based feature selection with network-based strategies. This integrative design ensures that candidate taxa are supported by both predictive modeling and ecological network context, thereby increasing confidence in their biological relevance. The main contributions of this study are as follows: 1) We present a novel Bayesian network-based algorithm, implemented in the R package `CMIMN`; 2) We propose a consensus network approach that integrates results from `CMIMN`, `SPIEC-EASI`, `SPRING`, and `SPARCC`, improving the reliability of inferred microbial associations; and 3) We develop a dual feature selection strategy that incorporates both machine learning outputs and network centrality metrics to identify key disease-associated taxa. Importantly, the methodology introduced here is broadly applicable and can be adapted for analyzing disease-related microbiome datasets across both agricultural and clinical domains.

## 2 Materials and methods

Figure 1 shows a graphical representation of our pipeline with four major panels. Each panel represents a specific step:

- **Data Preparation:** Microbiome abundance data is filtered to retain operational taxonomic units (OTUs) present in at least 15 samples, ensuring that low-prevalence taxa do not introduce noise. The resulting filtered data matrix serves as the foundation for downstream analyses.
- **Constructing Microbiome Networks:** Microbiome networks are constructed using four methods—`SE_glasso`, `SPRING`, `SPARCC`, and the proposed `CMIMN`. These networks represent microbial taxa as nodes and their co-occurrence relationships as edges. To enhance reliability, a consensus microbiome network is constructed by integrating results from all four methods. Edge weights indicate the level of agreement among methods, with a weight of 4 representing relationships confirmed by all methods and 0 indicating no agreement.
- **Feature Selection (machine learning (ML)):** The filtered data is normalized using three methods (CLR, log, and TSS) and subjected to different ML-based feature selection methods. Each method assigns a “TOTAL” score to each OTU based on how frequently it is selected as important across the seven strategies. OTUs with high “TOTAL” scores are considered key features for further analysis.
- **Feature Selection (network-based):** Networks are separately constructed for ‘scab-infected’ and ‘clean tuber’ samples using `SE_glasso`, `SPRING`, `SPARCC`, and `CMIMN`. Topological features are computed for each node in the networks. Two distinct strategies—differential centrality analysis and weighted scoring of OTUs—are applied to identify important OTUs based on network structure.

**Figure 1:**
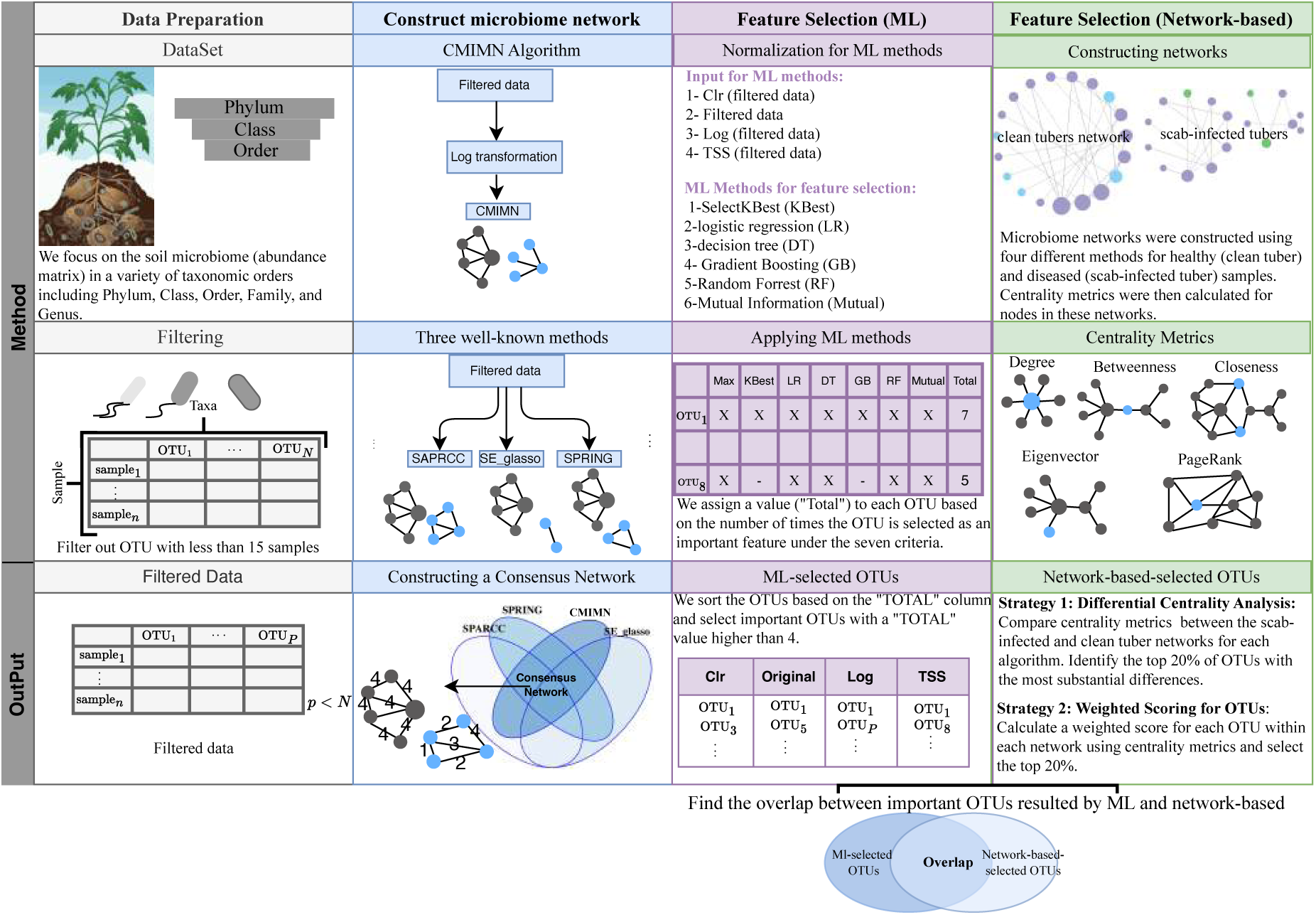
Workflow of the microbiome analysis pipeline for identifying key microbial drivers of disease resistance. Using potato common scab as an example, this pipeline consists of five main steps and is generally applicable to any microbiome dataset: (1) Data Preparation – Raw microbiome data is preprocessed to retain OTUs present in at least 15 samples, ensuring that low-prevalence taxa do not introduce noise. The resulting filtered data matrix serves as the foundation for downstream analysis, focusing on taxonomic levels such as Phylum, Class, and Order. (2) Construct microbiome network – Microbiome networks are constructed using four inference methods: `SE_glasso`, `SPRING`, `SPARCC`, and the proposed `CMIMN`. These networks represent microbial taxa as nodes and their interactions as edges. To improve reliability, a consensus microbiome network is constructed by integrating results from all four methods. Edge weights indicate the level of agreement among methods, with a weight of 4 representing relationships confirmed by all methods and 0 indicating no agreement. (3) Feature Selection (ML) – The filtered data undergoes normalization using three methods (CLR, log, and TSS) before applying ML-based feature selection methods. A “TOTAL” score is assigned to each OTU based on its selection frequency across ML methods, identifying key taxa strongly associated with disease outcomes. (4) Feature Selection (Network-Based) – Microbiome networks are separately constructed for ‘scab-infected’ and ‘clean tuber’ samples. Two strategies are applied to identify key OTUs based on network structure: (i) Differential Centrality Analysis, and (ii) Weighted Scoring of OTUs. Final OTU Selection: Identifying Overlap– The last step identifies the overlap between OTUs selected by ML-based and network-based approaches, ensuring robust and reliable feature selection for downstream microbiome analysis.

The Venn diagram at the bottom illustrates the overlap between OTUs identified by ML-based methods and network-based feature selection strategies. This overlap highlights the robustness of the selected OTUs as key contributors to disease resistance. This workflow integrates statistical rigor and biological relevance, ensuring that the identified OTUs are reliable targets for further investigation and potential microbial intervention strategies.

### 2.1 Data preparation

In this study, we focus on the soil microbiome (matrix of abundances) in a variety of taxonomic orders, including Phylum, Class, and Order from soil samples acquired from potato fields in Wisconsin and Minnesota. We concentrated on these three taxonomic levels due to their balance of interpretability and feature dimensionality, allowing for a meaningful analysis of microbial community structure and its association with disease.

The dataset consists of microbial community data of pre-planting soils and the corresponding disease levels in the plants at harvest. Overall, we collected 256 soil samples, 108 of which were taken from 36 commercial fields in Minnesota, and 148 of which were taken from 50 fields in Wisconsin. This extensive dataset provides a comprehensive representation of soil microbial communities across two major potato-growing regions in the Upper Midwest.

DNA was extracted from the the pre-planting soils, and analyzed for microbial community data following the method in [18]. Bacterial and fungal DNA was sequenced to capture a diverse range of microbial taxa, enabling us to investigate their interactions and potential roles in disease resistance or susceptibility. At harvest, potatoes were hand-harvested from a one-meter hill (usually 3-4 plants) at each sampling location. Tubers from one plant were visually evaluated for the presence of pitted scab lesions: which is a sign for serious common scab disease, as these tubers would be excluded from marketable yield. This binary disease label, 0 for healthy and 1 for diseased, serves as the target variable in our analyses, linking microbial community features to agricultural outcomes.

The input data is a matrix with non-negative read counts that were generated by a sequencing procedure, filtered out so that we only include OTUs that appear in at least 15 samples [18]. This filtering ensures a focus on microbial taxa with sufficient prevalence to contribute meaningfully to statistical and network-based analyses, reducing noise from rare taxa. Table S1 displays the number of features (OTUs) before and after filtering for different taxonomic levels. To enhance reproducibility and facilitate further research, the raw sequencing data and preprocessing scripts are available upon request or through the project repository.

### 2.2 Constructing microbiome networks

In the “Conditional mutual information algorithm for constructing microbiome networks (`CMIMN`)” section, we introduce a novel Bayesian Network-based approach, `CMIMN`, which leverages Conditional Mutual Information (CMI) to infer microbial associations in a more robust and scalable manner. However, despite advancements in microbiome network inference, no single method consistently produces reliable results due to inherent differences in statistical assumptions, data preprocessing, and sparsity constraints. Each inference method captures different aspects of microbial interactions, often leading to inconsistencies across constructed networks. Therefore, in the “Consensus network construction” section, we present a consensus microbiome network that integrates results from multiple inference methods to enhance reliability and mitigate method-specific biases.

#### 2.2.1 Conditional mutual information algorithm for constructing microbiome networks (CMIMN)

We outline the methodology behind the `CMIMN` algorithm, a novel approach for constructing microbiome networks. First, we introduce the foundational concepts of Mutual Information (MI) and Conditional Mutual Information (CMI), which are key components of the `CMIMN` framework. Next, we provide an overview of Bayesian Networks, their structure, and their applicability to microbiome research. Finally, we describe the detailed steps of the CMIMN algorithm, highlighting its dynamic thresholding and order-independent features that address the unique challenges of microbiome datasets.

##### Mutual information and conditional mutual information

MI and CMI are proven to be effective for detecting relationships between variables due to their capability to measure nonlinear dependencies [20]. MI and CMI between the variables *X* and *Y*, given the vector of variables **Z**, are defined as follows [21]:

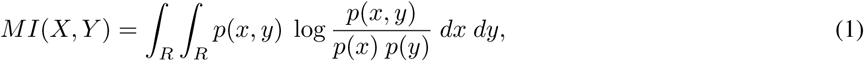

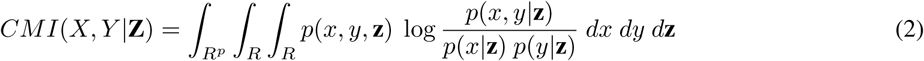

where *p* is the dimension of vector **Z** and *p*(*x, y*), *p*(*x*) and *p*(*y*) represent the joint distribution of *X* and *Y*, marginal distribution of *X*, marginal distribution of *Y*, respectively. *p*(*x, y,* **z**), *p*(*x, y|***z**), *p*(*x|***z**) and *p*(*y|***z**) indicate joint distribution of *X*, *Y* and **Z**, the conditional density distribution of *X* and *Y* given **Z**, the conditional density distribution of *X* given **Z** and the conditional density distribution of *Y* given **Z**, respectively. Under the assumption that data follows a Gaussian distribution, MI for two continuous variables *X* and *Y* can be calculated as [16, 22, 23]:

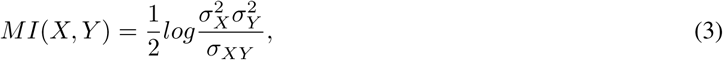

where 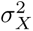, 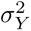 and *σ_XY_* indicate the variance of *X*, the variance of *Y* and the covariance between *X* and *Y*, respectively. When *X* and *Y* are independent, then *MI*(*X, Y*) = 0. Similarly, *CMI*(*X, Y |***Z**) is defined as:

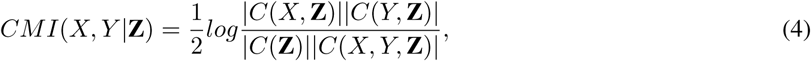

where C is the covariance matrix and *|.|* is the determinant of matrix C. C(X,Y) and C(X,Y, **Z**) denote the covariance matrix of variables *X* and *Y* and variables X, Y, and **Z**, respectively. When *X* and *Y* are conditionally independent given **Z**, then *CMI*(*X, Y |***Z**) = 0. These measures form the backbone of many network inference methods, including Bayesian Networks (BNs), which are particularly suited for capturing complex dependencies in microbiome datasets. Below, we provide an overview of BNs and their applications.

##### Overview of Bayesian networks

BNs are probabilistic graphical models that represent complex relationships among variables using directed acyclic graphs (DAGs). Each node in a BN represents a random variable, while the directed edges capture conditional dependencies between them. BNs have been extensively applied in various biological network analyses, such as gene regulatory networks [24–26], but their use in microbiome research remains limited. There are three main approaches for learning the structure of BNs: Constraint-Based Methods: These are based on conditional independence tests to infer the network structure [22, 24, 25, 27–31]. Score-Based Methods: These involve optimizing a scoring function to search among candidate network structures [32–34]. Hybrid Methods: These combine elements of both constraint-based and score-based approaches to leverage their respective strengths [16,35–39]. Among these, the PC algorithm and its derivatives, such as Fast Causal Inference, Really Fast Causal Inference, and `PCA-CMI` [22, 29, 30, 40–42], are prominent constraint-based methods. Despite their widespread use, these methods have notable limitations: 1-Order Dependence: The results can vary depending on the sequence in which the nodes are processed [43]. 2-Static Threshold Dependency: Using fixed thresholds for conditional independence tests often leads to false positives or false negatives, reducing the reliability of inferred networks [24].

##### CMIMN algorithm

We propose the `CMIMN` algorithm to overcome the challenges posed by microbiome data, providing an order-independent, dynamically threshold, and sparsity-controlled framework for microbiome network construction:

###### 1) Order independence

Traditional PC-based methods, such as `PCA-CMI`, are susceptible to order dependence, where the sequence in which nodes are processed affects the inferred network structure. This occurs because, in these methods, the tests for conditional independence and edge removal are performed simultaneously during each iteration, making the results highly sensitive to the order of node traversal. In contrast, `CMIMN` eliminates this dependency by decoupling these steps. Specifically, for each step of the algorithm, `CMIMN` begins by fixing the set of potential separators for every edge (*X, Y*). This set is determined as the intersection of the neighbors of *X* and *Y* in the current graph. By defining the separators upfront, the algorithm ensures that all configurations are consistently evaluated, regardless of the order in which nodes or edges are processed. Once the potential separators are fixed, the algorithm proceeds to calculate the independence measures (e.g., CMI) for each edge using the predefined separator sets. Edges that fail the independence test, based on the dynamically determined thresholds, are then removed. This sequential separation of tasks — fixing separators, calculating independence measures, and then removing edges — ensures that the outcome of each step is independent of the traversal order of the nodes.

###### 2) Dynamic thresholds

In traditional PC-based methods an edge between two nodes is removed if the independence measure (e.g., MI or CMI) falls below a predefined static threshold, *θ*, usually 0.05. However, this fixed-threshold approach is inherently rigid and can lead to significant issues. Specifically, static thresholds are often poorly calibrated to the scale and variability of different datasets, resulting in false positives (retaining spurious edges) or false negatives (removing meaningful edges). To overcome these limitations, `CMIMN` employs a quantile-based dynamic threshold approach. Instead of using a single, static threshold, thresholds are adaptively determined based on the statistical properties of the dataset. For example, in each iteration, thresholds are set using specific quantiles (e.g., the 70th percentile) of the computed MI or CMI values. This ensures that the threshold dynamically adjusts to the distribution of independence measures, accommodating variability in the data’s scale, density, and characteristics.

###### 3) Sparsity control

Network sparsity is a crucial factor in microbiome studies, as overly dense networks can obscure biologically meaningful interactions, while overly sparse networks may omit key relationships. In traditional PC-based methods, selecting a single static threshold does not allow for precise control over network sparsity. However, CMIMN addresses this challenge by offering precise control over network sparsity through quantile-based criteria for edge removal. By dynamically tuning the sparsity threshold, researchers can specify the desired percentage of edges to retain, ensuring that the resulting network retains significant edges representing meaningful microbial interactions while reducing noise and redundancy. The steps of the `CMIMN` algorithm are outlined below:

*Step 0: Initialization:* Generate a complete network with the number of nodes equal to the number of taxa.
*Step 1: Calculate MI of Order 0:* Compute MI values for each pair of nodes.
*Step 2: Remove Edges:* Remove edges for which MI values are smaller than *θ*_1_, the threshold for the MI test. The resulting network at this stage is denoted by *S*_0_.
*Step 3: Calculate CMI of Order 1:* Calculate *CMI*(*X, Y* |*Z*) where *Z* belongs to the set *V_XY_* = *ADJ*(*X*) ∩ *ADJ*(*Y*) in *S*_0_. Here, *ADJ*(*X*) represents the set of nodes that are adjacent to *X*. We consider ‘paths of length 2 between *X* and *Y*’ to mean that *Z* is a common neighbor of both *X* and *Y*. Thus, a path from *X* to *Y* via *Z* consists of two edges: one connecting *X* to *Z* and another connecting *Z* to *Y*. This configuration is used to assess the indirect interactions between *X* and *Y* mediated by *Z*, focusing on how *Z* influences the dependency between *X* and *Y*.
*Step 4: Remove Edges:* Define *CMI*_70_ (*X, Y* |*Z*) as the 70th percentile of all *CMI*(*X, Y* |*Z*) values. If *CMI*_70_ (*X, Y* |*Z*) *< θ*_2_ (the threshold for the CMI test), remove the edge between *X* and *Y*. The resulting skeleton at this stage is denoted by *S*_1_.
*Final Outcome:* The resulting network *S*_1_ is a fully undirected skeleton.

The primary challenge in using BN methods to infer microbiome networks lies in the normalization of count datasets for the BN algorithm. To address this challenge, we apply a logarithmic transformation to the count data to stabilize variance, reduce skewness, and address compositional constraints inherent in microbiome datasets. This transformation ensures that MI and CMI operate on a continuous, normalized scale, enhancing their reliability. Without this step, applying MI or CMI directly to raw counts would yield biased results due to the data’s non-normalized and highly variable nature. The `CMIMN` algorithm is implemented in R and publicly available in https://github.com/solislemuslab/CMIMN.

#### 2.2.2 Constructing a consensus network

We apply three state-of-the-art network methods on the soil microbiome data: 1) SParse InversE Covariance Estimation for Ecological Association Inference (`SPIEC-EASI`) method [44], using the graphical lasso option (referred to as `SE_glasso`); 2) `SPRING`: Semi-Parametric Rank-based approach for Inference in Graphical Models [45], and 3) `SPARCC`: Sparse Correlations for Compositional Data [46]. The input of these methods is an abundance matrix and the output is undirected networks in which nodes represent OTUs and edges corresponds to interactions between them.

First, `SE_glasso` [44] estimates sparse inverse covariance matrices to infer ecological associations in microbial communities. The approach is designed to address the challenges of compositional data and high dimensionality commonly encountered in microbiome studies. By accurately modeling microbial interactions and leveraging graphical lasso regularization, `SE_glasso` uncovers significant ecological relationships between different taxa.

Second, the `SPRING` [45] algorithm is a powerful method for inferring associations in complex biological networks that combines the advantages of both parametric and non-parametric approaches to construct a co-occurrence network from abundance data, commonly encountered in microbiome studies. By transforming the abundance values into ranks and utilizing rank-based statistical tests, `SPRING` overcomes the challenges of compositional data and improves robustness against outliers and extreme values. This algorithm effectively identifies significant interactions between different taxa, providing valuable insights into the underlying structure and ecological relationships within microbial communities.

Third, `SPARCC` [46] is a powerful algorithm used to analyze microbial communities and infer associations between different taxa. Specifically designed for compositional data, which represents the relative abundances of microbial taxa, `SPARCC` addresses the challenges of dealing with non-negative and constrained data. By regularizing the correlation matrix through a bootstrap procedure and using sparsity-inducing techniques, `SPARCC` efficiently estimates sparse correlations between taxa, revealing significant co-occurrence patterns and potential ecological interactions within the microbial community. The algorithm’s ability to handle compositional data makes it a valuable tool for investigating complex microbial ecosystems and unraveling the underlying relationships between taxa.

Building reliable microbiome networks is difficult because different algorithms produce varying results. Each method has its own strengths and weaknesses, which can lead to inconsistent interpretations of microbial interactions. To solve this, we combine the results from four methods to create a consensus network: `CMIMN`, `SE_glasso`, `SPRING`, and `SPARCC`. The consensus network is represented as a weighted adjacency matrix, where each edge (connection between two nodes) is assigned a weight value ranging from 0 (not identified by any algorithm) to 4 (identified by all four algorithms). This weight reflects the level of agreement among the algorithms regarding the presence of the edge. Indeed, the consensus network is constructed by first generating individual networks using each of the four methods. These networks are then overlaid, and the weight of each edge in the consensus network is computed as the sum of the binary indicators (presence/absence) of that edge across all four networks. The resulting weighted network not only highlights the most reliable edges (with higher weights) but also provides a comprehensive representation of microbial interactions. Among the advantages of the Weighted Consensus Network, we can highlight the *robustness* as integrating multiple algorithms reduces the impact of biases or errors associated with any single method; *stability* as the weighted approach provides a holistic view of microbial interactions, capturing edges that are consistently supported across algorithms; *interpretability* as the weight values offer a straightforward measure of edge confidence, allowing researchers to focus on highly reliable interactions for downstream analysis, and *sparsity Control* as selecting different threshold values for edge weights (e.g., retaining only edges with weights *≥* 1, *≥* 2, *≥* 3, or = 4) can control the sparsity of the network to match their analytical goals. For instance, lower thresholds (e.g., *≥* 1) result in denser networks that include more potential interactions, while higher thresholds (e.g., = 4) produce sparser networks focused only on the most reliable interactions identified by all algorithms. This weighted consensus network serves as a stable foundation for exploring microbiome interactions and identifying key microbial taxa and their relationships within complex ecosystems. By incorporating agreement across multiple methods, it offers a more reliable and nuanced perspective on the underlying microbial community structure.

### 2.3 Multi-method approach to identify key microbial drivers of disease resistance

Feature selection is a crucial step in data analysis, involving the identification of significant features or covariates that possess high predictive power. In the context of high-dimensional data, such as microbiome datasets, feature selection becomes indispensable to extract relevant information and reduce computational complexity. In particular, when studying diseases, it becomes imperative to identify important OTUs that are strongly associated with the disease’s onset or progression. By identifying these key OTUs, we gain essential insights into potential driver pathogens or beneficial microbes. Subsequently, controlling the abundance or activity of these crucial OTUs can pave the way for novel disease interventions and management strategies, opening up avenues for precision medicine and tailored therapies. To identify key OTUs associated with disease outcomes, we employ a two-pronged feature selection approach: (1) Using machine learning-based methods, and (2) Using network-based methods. By integrating these complementary approaches, we ensure a comprehensive and biologically meaningful selection of microbial taxa associated with disease outcomes.

#### 2.3.1 Using machine learning-based methods

To identify important OTUs, we first normalize the filtered microbiome data using four different transformations: centered log-ratio (CLR), raw filtered data, logarithmic transformation (log), and total sum scaling (TSS). These normalization methods help account for compositional constraints and improve the reliability of machine learning (ML)-based feature selection.

We then apply all ML-based feature selection strategies, implemented in the `scikit-learn` library [47] in Python: (1) “SelectKBest” method, which selects features based on the *k* highest analysis of variance F-value scores, (2) Selection of the top *k* features based on the mutual information statistic, (3) Recursive Feature Elimination (RFE) with logistic regression, (4) RFE with a decision tree, (5) RFE with gradient boosting, and (6) RFE with Random Forest (RF). Additionally, we introduce a 7th method that includes OTUs in the model if their maximum value falls within the top 30% of the dataset. Running all seven strategies, we assign a value (“TOTAL”) to each OTU based on the number of times the OTU is selected as an important feature under the seven criteria. Specifically, an OTU that is selected as important by all seven strategies will have “TOTAL” value of 7. Subsequently, we sort the OTUs based on the “TOTAL” column and select important OTUs with a “TOTAL” value higher than a defined threshold. These approaches collectively provide valuable insights into the most influential OTUs in the context of our feature selection analysis, allowing us to make informed decisions and draw meaningful conclusions in the subsequent stages of our study.

#### 2.3.2 Using network-based methods

While machine learning-based methods identify statistically relevant OTUs, they do not capture microbial interactions that may play a crucial role in disease resistance. To address this limitation, we employ a network-based feature selection strategy that compares microbial co-occurrence patterns between diseased and healthy samples. We construct microbial interaction networks using four well-established methods (`1-SPARCC`, `2-SE_glasso`, `3-SPRING`, and `CMIMN`) based on samples from two classes: one representing samples without the disease (‘clean tubers’) and the other with the disease (‘scab-infected tubers’). We then apply two complementary network-based feature selection strategies:

##### Strategy 1: differential centrality analysis

This approach analyzes five centrality metrics for each OTU: 1-Degree (connectivity within the network), 2-Betweenness (importance in connecting other taxa), 3-Closeness (proximity to all other taxa), 4-Eigenvector Centrality (influence based on connected neighbors), and 5-PageRank (importance based on link structure). We rank OTUs based on the **difference** in their centrality measures between the ‘clean tubers’ and ‘scab-infected tubers’ networks. The top 20% of OTUs showing the most significant variations are selected as key taxa. The final taxa are considered important if they are selected by **all** four network inference methods.

##### Strategy 2: weighted scoring of OTUs based on network topology

This method assigns a weighted score to each OTU based on its network properties using the following formula:

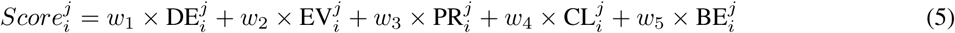

where, *i* denotes the OTU being evaluated and *j* represents the network inference method used for constructing the microbiome network. Here, DE, EV, PR, CL, and BE represent Degree, Eigenvector, PageRank, Closeness, and Betweenness centrality measures, respectively. The weights assigned to the centrality measures reflect their relative importance in capturing biologically meaningful insights from the network structure. To ensure that the OTUs identified as significant have greater overlap with Strategy 1, we set the weights as follows: *w*_1_ = 0.1, *w*_2_ = 0.1, *w*_3_ = 0.1, *w*_4_ = 0.2, *w*_5_ = 0.5, with Betweenness (50%) given the highest weight due to its role in structuring the microbial community. The final top 20% of OTUs with the highest scores are considered key players in microbial interactions related to disease resistance.

##### Unifying machine learning and network-based approaches for reliable microbiome feature selection

To improve the reliability of microbiome feature selection, we integrate both machine learning and network-based strategies. Specifically, we evaluate OTUs identified by two different network-based methods with those selected through multiple machine learning algorithms. The final set of selected OTUs consists of taxa consistently prioritized across these complementary approaches, thereby increasing confidence in their biological relevance. This integrative strategy combines the predictive power of machine learning with the structural insights derived from microbial interaction networks, resulting in a robust and interpretable set of microbial features. The selected OTUs represent strong candidates for microbial drivers of disease resistance and may inform the development of microbiome-targeted interventions aimed at enhancing crop resilience and promoting sustainable agricultural practices.

## 3 Results

### 3.1 Robustness study of different algorithms for learning the microbiome network

In order to assess the robustness of CMIMN algorithm in learning the microbiome network, we conducted a comprehensive analysis. In the initial step, we constructed the network utilizing the entirety of all samples. Subsequently, we performed a critical evaluation by randomly selecting 70% of the samples and generating 50 distinct datasets derived exclusively from this subset. On each of these 50 datasets, we executed the algorithm independently to construct separate networks. We compare the networks constructed from each of the 50 generated datasets to the corresponding network generated using the complete set of samples, for each method based on F-score. This comparative evaluation assesses how consistently each algorithm reconstructs microbial associations across different generated of the data. The F-score is a widely used metric that balances precision and recall, providing a single measure of a method’s accuracy in detecting true microbial interactions while minimizing false positives and false negatives. A higher F-score indicates better network reconstruction accuracy and reliability. To visually represent these comparisons, we created box plots for the F-score values obtained from each iteration. This allowed me to not only assess the overall performance but also identify any potential variations in performance across different taxonomic levels, including Phylum, Class, and Order. Furthermore, we extended this rigorous evaluation to encompass various network construction methods, including CMIMN, SPRING, SE_glasso, and SPARCC. The comparison was conducted at each taxonomic order, resulting in a comprehensive assessment of the method’s robustness under different conditions. Figure S1 presents the box plots showcasing the F-scores resulting from the application of different methods at varying taxonomic orders. This visualization provides a clear and insightful representation of the method’s performance across different scenarios, offering valuable insights into its reliability and effectiveness. Performance comparisons were made across different taxonomic levels, including Phylum, Class, and Order. Our algorithm, CMIMN, exhibits superior performance as indicated by the narrower range of box plots in all three taxonomic levels, demonstrating its robustness. Notably, among all algorithms, SE_glasso shows the least favorable results for Class level.

### 3.2 Minimal Overlap Across Network Inference Methods Highlights the Need for Using a Consensus Approach

We constructed microbiome networks using four different inference methods: (SE_glasso, SPRING, SPARCC, and CMIMN) at the Phylum, Class, and Order levels. Despite using the same dataset, the resulting networks exhibited minimal overlap, highlighting the high variability in microbial interaction patterns inferred by different methods.

Figure 2 shows the Venn diagrams of common edges inferred by different methods at different taxonomic levels: only 24 common edges at the Phylum level, 80 at the Class level, and 522 at the Order level. These results indicate that network structures can vary significantly depending on the inference method used, which raises concerns about the reliability of conclusions drawn from any single approach.

**Figure 2:**
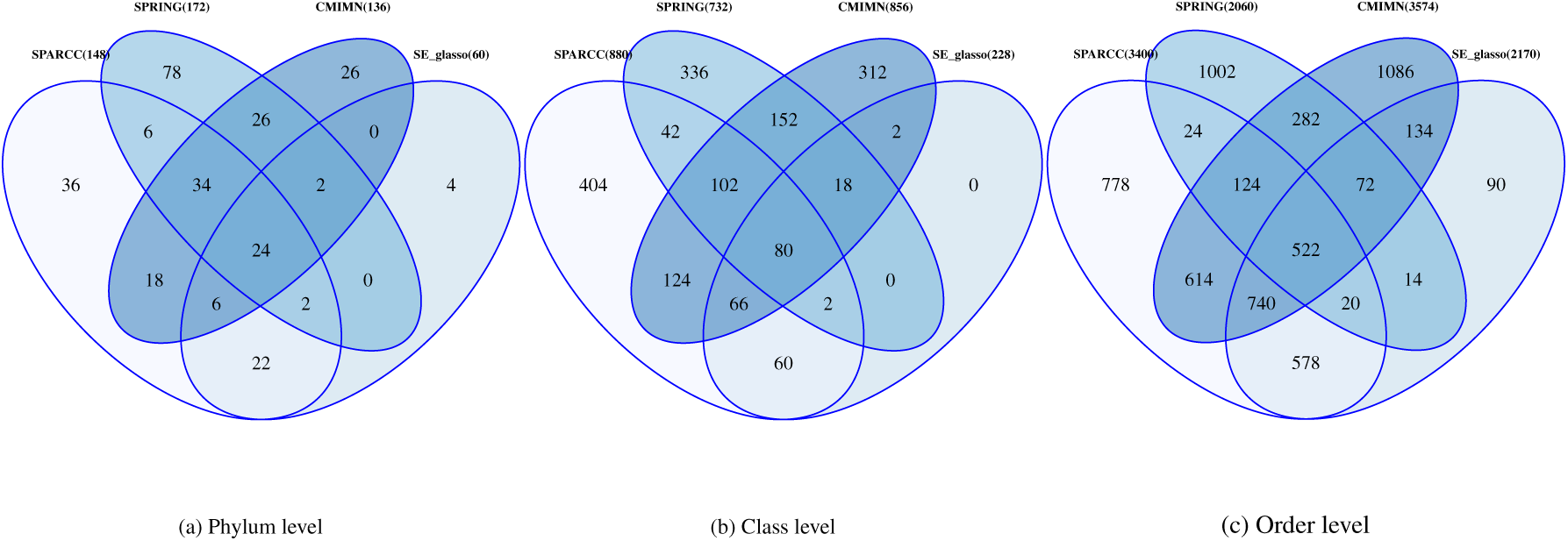
Venn diagrams illustrating the overlap of common edges in microbiome networks constructed using four different inference methods (`SE_glasso`, `SPRING`, `SPARCC`, and `CMIMN`) at different taxonomic levels based on all samples. (a) Phylum level: 24 common edges among all methods. (b) Class level: 80 common edges. (c) Order level: 522 common edges.

Table S2, S3, and S4 present network metrics for the four different methods at the Phylum, Class, and Order levels. The substantial differences in network topology metrics underscore the inherent differences in algorithmic assumptions and their impact on inferred microbial interactions. This variation highlights the importance of employing a consensus-based approach to enhance network robustness, reduce algorithm-specific biases, and improve biological interpretability.

Table S5, S6, and S7 present the top nodes in microbial networks based on topological measures at the Phylum, Class, and Order levels. These networks were constructed using `SE_glasso`, `SPRING`, `SPARCC`, and `CMIMN` methods, and the analysis encompasses data from all samples, without distinguishing between diseased and healthy conditions.

Interestingly, certain OTUs were consistently identified as highly connected nodes across all four network construction methods, reinforcing their biological significance. According to these tables, *Acidobacteriae* was identified as important at the Class level by all algorithms and network metrics, while *C0119* was consistently identified at the Order level. The recurrence of these taxa across multiple network inference methods suggests their potential ecological importance and role in microbial community stability.

### 3.3 Consensus microbiome network: enhancing reliability through integration

Figures 3, S2, and S3 visualize the microbiome networks at the Phylum, Class, and Order levels respectively. Part (a) of each figure represents the ‘clean tubers’ network, which is characterized by a denser and more connected microbial community, with more diverse interactions. Part (b) shows the ‘scab-infected tubers’ network, highlighting nodes and edges unique to diseased samples. Part (c) illustrates the common interactions between the two networks. Nodes represent OTUs and are color-coded: Purple: Common OTUs shared between ‘clean tubers’ and ‘scab-infected tubers’ networks. Blue: OTUs unique to the ‘clean tubers’ network. Green: OTUs unique to the ‘scab-infected tubers’ network. Node size reflects connectivity (degree), while edges are distinguished as dashed (confirmed by three methods) or solid (confirmed by all four methods). At the Class and Order levels, due to the density of edges, only solid edges (confirmed by all four methods) are reported for clarity.

**Figure 3:**
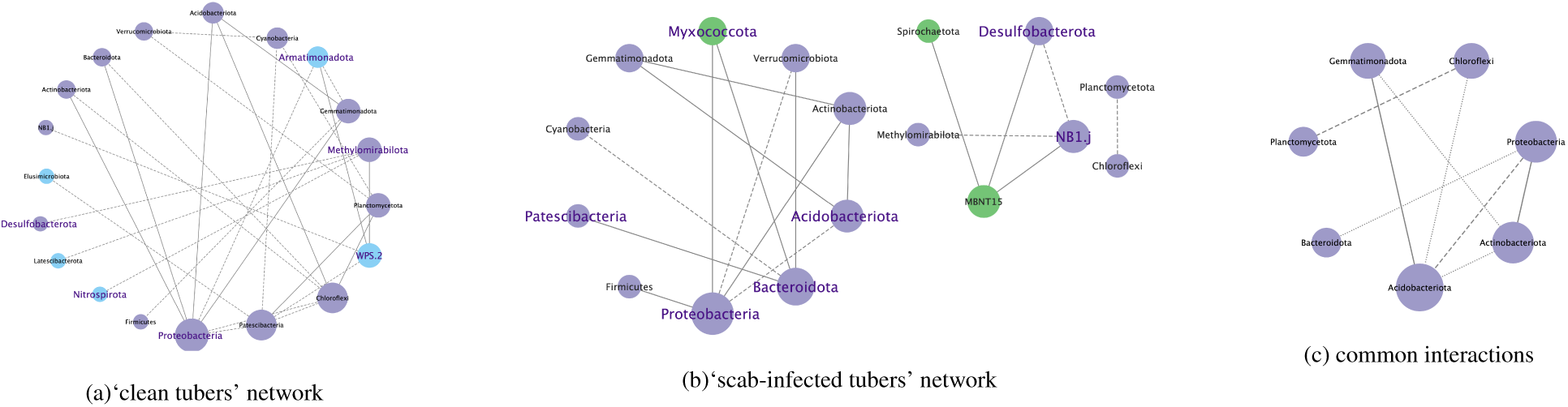
The microbiome network at the Phylum taxonomic level. Part (a) represents the ‘clean tubers’ network, part (b) displays the ‘scab-infected tubers’ network, and part (c) shows the common interactions between them. Nodes represent Operational Taxonomic Units (OTUs) and are color-coded: purple for common OTUs shared between ‘clean tubers’ and ‘scab-infected tubers’ networks, blue for OTUs unique to the ‘clean tubers’ network, and green for OTUs unique to the ‘scab-infected tubers’ network. Node size indicates their degree of connectivity. Edges are categorized as dashed lines (confirmed by three methods) or solid lines (confirmed by all four methods).

In both ‘clean tubers’ and ‘scab-infected tubers’ microbiome networks, we identified microbial associations consistently supported by all four network inference methods. Many of these associations have also been independently reported as ecologically meaningful in soil ecosystems. The intersections of microbial associations between the clean tubers’ and scab-infected tubers’ networks are summarized in Tables S8, S9, and S10, corresponding to the Phylum, Class, and Order levels, respectively. Each table includes three columns: the first lists associations unique to the ‘clean tubers’ network, the second shows associations exclusive to the ‘scab-infected tubers’ network, and the third highlights shared associations—those present in both networks—representing conserved or stable microbial interactions across conditions.

Due to the large number of associations observed at the Class and Order levels, we report the complete set of edges confirmed by all four methods in Supplementary Tables S9 and S10. Here, we focus on selected Phylum-level associations that are most frequently supported by existing literature. In the Table S8 for ‘clean tubers’ network, we observed the interaction between *Planctomycetota–Patescibacteria*, which likely reflects syntrophic or symbiotic interactions, as both phyla possess reduced genomes and are known to co-occur in structured soil aggregates [48]. The edge between *Methylomirabilota–WPS-2* may indicate shared adaptation to oligotrophic or co-contaminated soil conditions, where both phyla are often involved in carbon and nitrogen cycling under stress [49]. Additionally, *Gemmatimonadota–Proteobacteria* and *Gemmatimonadota–Acidobacteriota* were found only in the clean network; these phyla are often associated with nutrient cycling and stable soil conditions, suggesting cooperative metabolic roles in healthy tuber-associated soils.

In the ‘scab-infected network, the *Actinobacteriota–Gemmatimonadota* interaction was unique and mirrors findings from boreal forest soils, where both phyla jointly contributed to the transformation of dissolved organic matter during freeze–thaw cycles, pointing to potential functional synergy under stress [50].

In both ‘clean tubers’ and ‘scab-infected networks, the edges *Actinobacteriota–Proteobacteria* and *Proteobacte-ria–Acidobacteriota* were consistently present, suggesting stable and ecologically relevant relationships across conditions. The co-occurrence of *Actinobacteriota and Proteobacteria* has been reported in sandy and layered soils, where both phyla are dominant and likely contribute to complementary roles in organic matter degradation and nutrient cycling [51]. Meanwhile, Proteobacteria and Acidobacteriota are among the most abundant phyla in forest and agricultural soils and are known to occupy distinct but co-existing niches—Proteobacteria favoring copiotrophic (nutrient-rich) environments and Acidobacteriota preferring oligotrophic, acidic soils—indicating a functional partitioning that supports broad microbial diversity and resilience [52].

### 3.4 Feature selection using a multi-method approach

#### 3.4.1 Machine learning-based feature selection

Tables S11, S12, and S13 summarize the results of machine learning (ML)-based feature selection at the Phylum, Class, and Order levels, respectively. Each table includes five columns: four corresponding to different data normalization strategies applied prior to ML analysis, and a fifth representing their intersection. The first column (ML_CLR) reports OTUs selected from count-filtered data normalized using the CLR transformation. The second column (ML_Original) shows OTUs selected from raw count-filtered data without transformation. The third column (ML_Log) includes OTUs identified from log-transformed data, and the fourth column (ML_TSS) presents selections based on data normalized using TSS. The final column (ML_Intersection) lists OTUs consistently identified as important across all four normalization methods, highlighting microbial taxa that are robust to normalization choice. According to these tables, there is a good overlap between different normalization methods.

#### 3.4.2 Network-Based Selection: Strategy 1 (Differential Centrality Analysis)

Tables S14, S15, and S16 provide lists of selected OTUs at the Phylum, Class, and Order levels, respectively, based on Strategy 1: Differential Centrality Analysis. OTUs were selected according to two criteria: (1) those exhibiting the largest differences in centrality values between ‘clean tubers’ and ‘scab-infected tubers’ networks, and (2) those consistently identified across all four network inference methods (`SE_glasso`, `SPRING`, `SPARCC`, and `CMIMN`), reinforcing their biological relevance. The first column of these tables shows the OTU name, while the second column (Features) indicates the centrality measure(s) responsible for the OTU’s selection. These taxa represent microbial features whose connectivity patterns consistently differ between ‘clean tubers’ and ‘scab-infected tubers’ networks, suggesting a potential role in disease dynamics.

#### 3.4.3 Network-Based Selection: Strategy 2 (Composite Scoring Approach)

First, we constructed two distinct microbiome networks: one for ‘clean tubers’ and one for ‘scab-infected tubers’. Networks were generated using four inference methods: `SE_glasso`, `SPRING`, `SPARCC`, and `CMIMN`.

Next, we applied a composite scoring system that integrates multiple centrality metrics into a single weighted score for each OTU, as defined in Equation (5). This score was calculated separately for each of the four network inference methods. The top 20% of OTUs with the highest scores were considered significant. To evaluate the agreement between network-based selection (Strategy 2) and ML-based selection, we examined the overlap between the top 20% highest-scoring OTUs and those selected by ML-based methods.

Tables S17, S18, and S19 summarize these overlaps at the Phylum, Class, and Order levels, respectively. In these tables, the left section corresponds to results from the ‘clean tubers’ network, while the right section presents the results from the ‘scab-infected tubers’ network. Each section contains five columns: The first column lists the OTUs selected by Strategy 2. Columns 2–5 indicate whether the same OTUs were also identified by ML methods under four normalization strategies—CLR transformation, untransformed (raw) data, log-transformation, and TSS normalization. A value of “1” in these columns denotes agreement between Strategy 2 and the corresponding ML method for that normalization, while “0” indicates the OTU was uniquely selected by the network-based approach. This design enables a direct comparison of method overlap under different data preprocessing conditions. Figures S4 and S5 provide a visual representation of these overlaps, illustrating the average agreement between ML-based and network-based selection methods under different conditions. For ‘clean tubers’, at the Phylum level, `CMIMN` and `SPARCC` demonstrated slightly better agreement with ML-based selection across different normalization strategies. At the Order level, agreement was highest overall, particularly under CLR and TSS normalization. For ‘scab-infected’ tubers, `CMIMN` generally exhibited higher agreement with ML-based selection at the Phylum level, while `SPARCC` showed better alignment at the Class level in certain cases. At the Order level, both `CMIMN` and `SPARCC` consistently achieved strong agreement with ML-based methods. These findings indicate that `CMIMN` and `SPARCC` consistently align more closely with ML-based feature selection methods, particularly at the Order level and under CLR and TSS normalization. This underscores their robustness and reliability in identifying important OTUs across different experimental conditions.

Tables S20, S21, and S22 present the overlap between Strategy 2 and the ML approaches at the Phylum, Class, and Order levels, focusing specifically on OTUs that were commonly identified in **both** the ‘clean tubers’ and ‘scab-infected tubers’ networks for each method. The left sections of these tables report overlaps for the `CMIMN` and `SE_glasso` methods. The first column lists the microbial taxa consistently identified across both networks using each respective method. Columns 2 through 5 indicate whether these taxa were also selected by ML methods under different normalization strategies. A value of “1” denotes agreement between the ML and network-based methods, while “0” indicates no overlap. Similarly, the right sections of the tables summarize the results for the `SPARCC` and `SPRING` methods, following the same structure.

Finally, Tables S23 and S24 illustrate the overlap among important OTUs identified by all four algorithms (`CMIMN`, `SPARCC`,`SE_glasso`,`SPRING`)for the ‘clean tubers’ and ‘scab-infected tubers’ networks, respectively, based on Strategy 2.

### 3.5 Overall OTUs identified by all methods as key drivers of disease

Table 1 summarizes the key OTUs identified through different selection strategies, including Machine Learning-based feature selection (ML), Network-Based Selection: Strategy 1 (Differential Centrality Analysis) (Strategy 1), and Network-Based Selection: Strategy 2 (Composite Scoring Approach) (Strategy 2), at the Phylum, Class, and Order levels.

**Table 1:** Important taxa at the Phylum, Class, and Order levels across three different strategies. The symbols indicate intersections between the levels: * for taxa appearing in ML, Strategy 1, and Strategy 2; ▪ for taxa appearing in ML and Strategy 1; *•* for taxa appearing in ML and Strategy 2; ▲ for taxa appearing in Strategy 1 and Strategy 2.

| Level | ML | Strategy 1 | Strategy 2 | Intersection |
| --- | --- | --- | --- | --- |
| **Phylum** | Firmicutes | Bacteroidota | Bacteroidota | ▲ Bacteroidota |
|  | Cyanobacteria | WPS.2 | WPS.2 | ▲ WPS.2 |
|  | Armatimonadota | Proteobacteria | Proteobacteria | ▲ Proteobacteria |
|  | NB1.j |  |  |  |
| **Class** | Bacilli | Desulfotobacteriia | Ktedonobacteria | ▲ Actinobacteria |
|  | Ktedonobacteria | Actinobacteria | Vicinamibacteria | ▲ AD3 |
|  | Cyanobacteriia | Syntrophobacteria | Actinobacteria | ● Bacilli |
|  | Saccharimonadia | AD3 | Gammaproteobacteria | ● Anaerolineae |
|  | Planctomycetes |  | Alphaproteobacteria | ● Ktedonobacteria |
|  | Ignavibacteria |  | Acidobacteriae |  |
|  | Dehalococcoidia |  | Anaerolineae |  |
|  | Anaerolineae |  | AD3 |  |
|  | MB.A2.108 |  | Blastocatellia |  |
|  | Chthononomadetes |  | Bacilli |  |
| **Order** | Saccharimonadales | C0119 | C0119 | * C0119 |
|  | Bacillales | Defluviicoccales | Rhizobiales | ■ Defluviicoccales |
|  | C0119 | Bacteroidales | Chitinophagales | ■ Bacteroidales |
|  | Subgroup.2 | Kryptoniales | Ktedonobacterales | ● Ktedonobacterales |
|  | Xanthomonadales | B12.WMSP1 | Microtrichales |  |
|  | Acidobacteriales | Desulfotobacteriales |  |  |
|  | Chloroplast |  |  |  |
|  | Alicyclobacillales |  |  |  |
|  | Paenibacillales |  |  |  |
|  | Acetobacterales |  |  |  |
|  | Pseudomonadales |  |  |  |
|  | Anaerolineales |  |  |  |
|  | Elsterales |  |  |  |
|  | Bacteroidales |  |  |  |
|  | Ktedonobacterales |  |  |  |

**At the Phylum level,** we found no overlap between the taxa selected by ML-based and network-based methods, highlighting the complementary nature of these approaches. To provide a comprehensive view, we report the most consistently selected taxa within each category.

*ML-based selected taxa: Firmicutes*, identified through ML-based feature selection, comprise 5.5% of the bacterial community and promote plant growth and disease suppression, especially *Bacillus* spp., which enhance root colonization and pathogen inhibition through antimicrobial metabolites [53]. *Cyanobacteria* (0.9%) also emerged as important ML-selected taxa, contributing to nitrogen fixation, biofilm formation, and soil structure improvement, benefiting microbial community stability [54]. Less abundant phyla such as *Armatimonadota* (0.2%) may also be directly related to the disease, as negative relationship between the abundance of those phyla and soil suppressive ability of scab has been observed in one of our studies (data unpublished). The precise implications of *NB1-j* (0.1%) in disease progression are still unclear, but its involvement in nitrogen cycling and interactions with microalgae suggest potential indirect influences [55].

*Network-based selected taxa (intersection of both strategies): Bacteroidota*, *WPS.2*, and *Proteobacteria* were consistently identified across both network-based strategies, indicating strong and robust association with disease status. *Bacteroidota* (5.5%) are involved in nutrient cycling and pathogen competition, both of which contribute to disease suppression [56]. *WPS.2*, though less prevalent (0.3%), showed a negative relationship with suppressive soil capacity in our prior (unpublished) observations. *Proteobacteria* represent a taxonomically diverse group containing both beneficial (e.g., *Rhizobium*) and pathogenic (e.g., *Pseudomonas syringae*, *Ralstonia solanacearum*) members, reflecting their complex role in disease ecology [57].

**At the Class level,** *Network-based intersection: Actinobacteria* and *AD3* were selected by both network-based strategies, indicating strong structural importance in microbial networks associated with disease. *Actinobacteria* are key contributors to soil suppressiveness against plant pathogens. Notably, non-pathogenic *Streptomyces spp.* produce antibiotics that inhibit soil-borne pathogens, including Streptomyces scabies, the causative agent of common scab disease [58]. Although less well characterized, Although less well-characterized, *AD3* was identified as a robust Class-level taxon across both network-based strategies. This group has been associated with degraded or polluted soils and reduced organic matter content, suggesting its presence may indicate shifts in microbial community structure linked to soil stress and disease vulnerability [59].

*ML and network-based Strategy 2 intersection: Bacilli*, *Anaerolineae*, and *Ktedonobacteria* were jointly identified by ML-based feature selection and network-based Strategy 2, suggesting these taxa are both predictive and structurally central in the disease-associated microbiome. *Bacilli* (notably *Bacillus* spp.) are widely recognized for their role in plant protection and disease suppression, particularly through Bacillus spp., which produce lipopeptides and hydrolytic enzymes that enhance root colonization and pathogen inhibition [60]. *Ktedonobacteria* exhibit complex morphologies and genomic features, leading to speculation that they may be a valuable microbial resource for novel compounds [61]. *Anaerolineae*, frequently found in low-oxygen soil habitats, play a crucial role in carbon degradation processes, including the breakdown of plant-derived compounds [62, 63]. This activity can modify the soil environment, potentially suppressing plant diseases through nutrient competition or the production of inhibitory substances.

**At the Order level,** *Confirmed by all ML methods and both network-based strategies: C0119* was the only taxon consistently identified by all machine learning models and both network-based strategies, highlighting its strong and stable association with disease-relevant microbial networks. Although taxonomically unclassified, recent studies have shown that *C0119* is a dominant order in biochar-amended soils, environments known to support improved microbial diversity, carbon cycling, and root-associated community stability [64]. Its consistent emergence across diverse analytical methods suggests an ecologically meaningful role in shaping disease-conducive or suppressive soil environments.

*Selected by ML and Network Strategy 1: Defluviicoccales*, while often linked to anaerobic degradation, have been observed in disease-prone soils, where they may contribute to microbial shifts influencing pathogen persistence [65]. *Bacteroidales* are involved in organic matter degradation and nutrient cycling. Some members have been associated with pathogen suppression via competitive exclusion and enhancement of soil nutrient availability, contributing indirectly to disease resistance [66].

*Selected by ML and Network Strategy 2: Ktedonobacterales* have been associated with disease suppression due to their potential for producing antimicrobial compounds and their metabolic similarity to antibiotic-producing Actinomycetes [61].

## Conclusion

This study introduces a comprehensive framework for robust microbiome network inference and the identification of disease-associated microbial taxa, specifically in the context of potato common scab. We developed CMIMN, a novel Bayesian network algorithm based on conditional mutual information, which exhibited superior robustness and interpretability across taxonomic levels. Recognizing the limitations of individual network inference methods, we integrated CMIMN with three widely used approaches—SPIEC-EASI, SPRING, and SPARCC—to construct consensus microbiome networks. These consensus networks captured biologically meaningful co-occurrence patterns while reducing algorithm-specific variability, thereby enhancing confidence in the inferred microbial interactions.

To identify taxa relevant to disease, we implemented a multi-method feature selection framework combining different machine learning algorithms with two network-based strategies. The machine learning component provided predictive stability by selecting features that consistently appeared across models, while the network-based methods leveraged centrality metrics and topological differences between networks to capture taxa important to microbial structure and community dynamics. This integrative approach enabled the detection of microbial taxa with both statistical significance and ecological relevance.

Our results revealed clear distinctions in microbial community structure between clean and scab-infected tubers. At the Phylum level, *Bacteroidota*, *WPS-2*, and *Proteobacteria* were identified through both network-based strategies, while *Firmicutes* and *Cyanobacteria* were highlighted by machine learning models. Interactions such as *Actinobacteriota–Proteobacteria* and *Planctomycetota–Patescibacteria* were found to be consistently supported by all four network inference methods and corroborated by existing soil studies, reinforcing their ecological relevance.

At the Class level, *Actinobacteria*, *AD3*, *Bacilli*, *Anaerolineae*, and *Ktedonobacteria* were identified by either multiple strategies or the intersection of both ML and network approaches. These classes are associated with key ecological functions such as carbon degradation, nutrient cycling, antimicrobial production, and disease suppression. At the Order level, *C0119* was the only taxon confirmed by all machine learning models and both network-based strategies, highlighting its potential as a robust indicator of disease status. Other important orders included *Bacteroidales*, *Defluviicoccales*, and *Ktedonobacterales*, identified by at least two independent methods.

The topological analysis of microbial networks further revealed differences in connectivity and interaction density between clean and diseased tuber microbiomes. Clean tuber networks exhibited higher overall connectivity, suggesting a more stable and cooperative microbial community. In contrast, disease-associated networks were more fragmented and featured shifts in taxa centrality, indicating structural reorganization in response to pathogen pressure. Several interactions identified in these networks—such as *Actinobacteriota–Gemmatimonadota* and *Methylomirabilota–WPS-2*—have also been observed in previous studies investigating soil response to stress or contamination.

The concept of disease-suppressive soil suggests fundamental differences in the microbiological environment between healthy and disease-conductive soils; meanwhile, soil microorganisms have been proposed as bioindicators of general soil health [67]. Because the microbiomes in this study were extracted from pre-planting soils of potato fields, those selected microbial features which distinct healthy and disease-conductive soils seem to exist long before disease emergency. Despite the variations driven by geography, management, and climate legacy in this large-scale survey, microbial signal was strong for those that tend to produce disease-free tubers. This suggests promising utility of soil microbiome in inferring indicators for soil health and predicting potato scab diseases.

Altogether, our integrative approach provides a scalable and interpretable framework for microbiome network analysis and biomarker discovery. By combining Bayesian inference, consensus-based network construction, and multi-method feature selection, we bridge predictive modeling with ecological insight. These findings not only improve our understanding of microbial community dynamics in disease contexts but also establish a foundation for microbiome-informed strategies in plant health management and sustainable agriculture. As a next step, our broader vision is to develop an interactive Shiny application that enables biologists to upload their microbiome and disease data to identify reliable taxa-disease associations and uncover robust co-occurrence relationships—making advanced analysis tools more accessible and actionable for the biological research community. In addition, our approach extracts biologically meaningful information by comparing network structures between clean tubers and scab-infected tubers: Strategy 1 focuses on differences in centrality measures across the two networks, while Strategy 2 analyzes each network independently to highlight key taxa. To further enhance this comparative analysis, we are interested in applying the Microbiome Network Alignment (MiNAA) algorithm [68], which aligns microbial networks across conditions, allowing us to extract deeper biological insights about shifts in microbial interactions associated with disease.

## Consent for publication

Not applicable.

## Availability of data and materials

The 16S and ITS amplicon sequencing data associated with this study are publicly available at the NCBI Short Read Archive under the BioProject PRJNA1135141. The R package of CMIMN and all R code for this paper are available in the github repository, https://github.com/solislemuslab/CMIMN.

## Authors’ contributions

CSL and RL developed the idea. RL and SS collected the data. RA led all statistical analyses from data preprocessing to fitting of machine learning models, as well as summarizing the results by the creation of figures. RA wrote the initial complete draft of the manuscript. SS, RL and CSL contributed in interpretations, editing, and revision of the manuscript. All authors read and approved the final manuscript.

## Supplementary Material

**Table 1:**
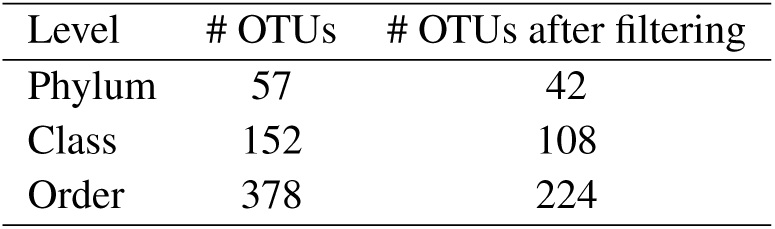
Taxonomic level (first column), number of OTUs in the original dataset (second column), and number of OTUs remaining after filtering out those that appear in fewer than 15 samples (third column).

| Level | # OTUs | # OTUs after filtering |
| --- | --- | --- |
| Phylum | 57 | 42 |
| Class | 152 | 108 |
| Order | 378 | 224 |

**Figure S1:**
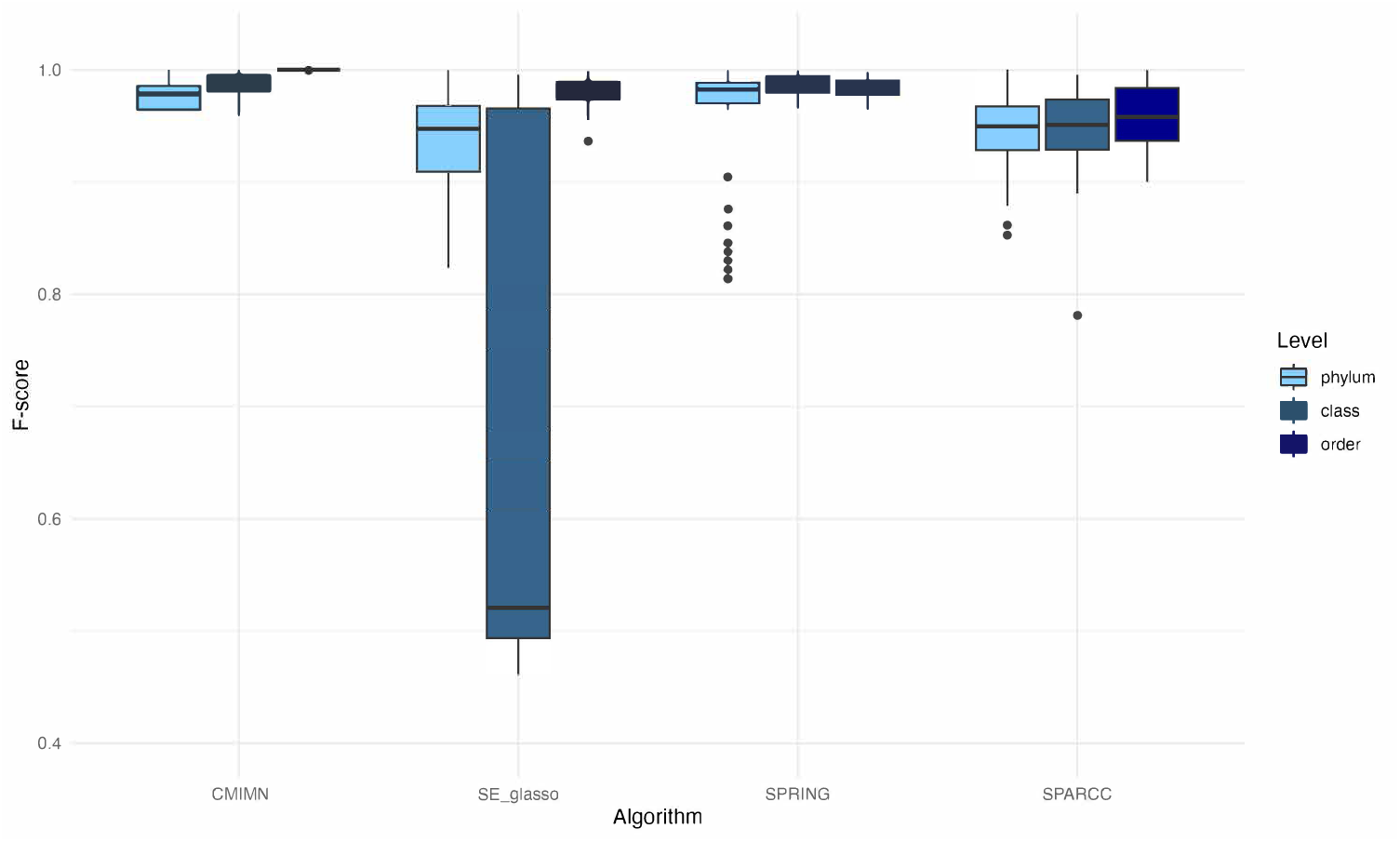
Box plots illustrating F-scores obtained from the robustness analysis of different microbiome network construction methods. The evaluation involved constructing networks from 50 distinct datasets, each generated by randomly selecting 70% of the samples from the full dataset. Performance was assessed across different taxonomic levels, including Phylum, Class, and Order. Four network inference algorithms—`CMIMN`, `SPRING`, `SE_glasso`, and `SPARCC`—were compared. Our results demonstrate that `CMIMN` exhibits superior robustness, as indicated by consistently higher and more stable F-scores across all taxonomic levels. Notably, `SE_glasso` shows the least favorable performance at the Class level, with greater variability and lower F-scores.

**Figure S2:**
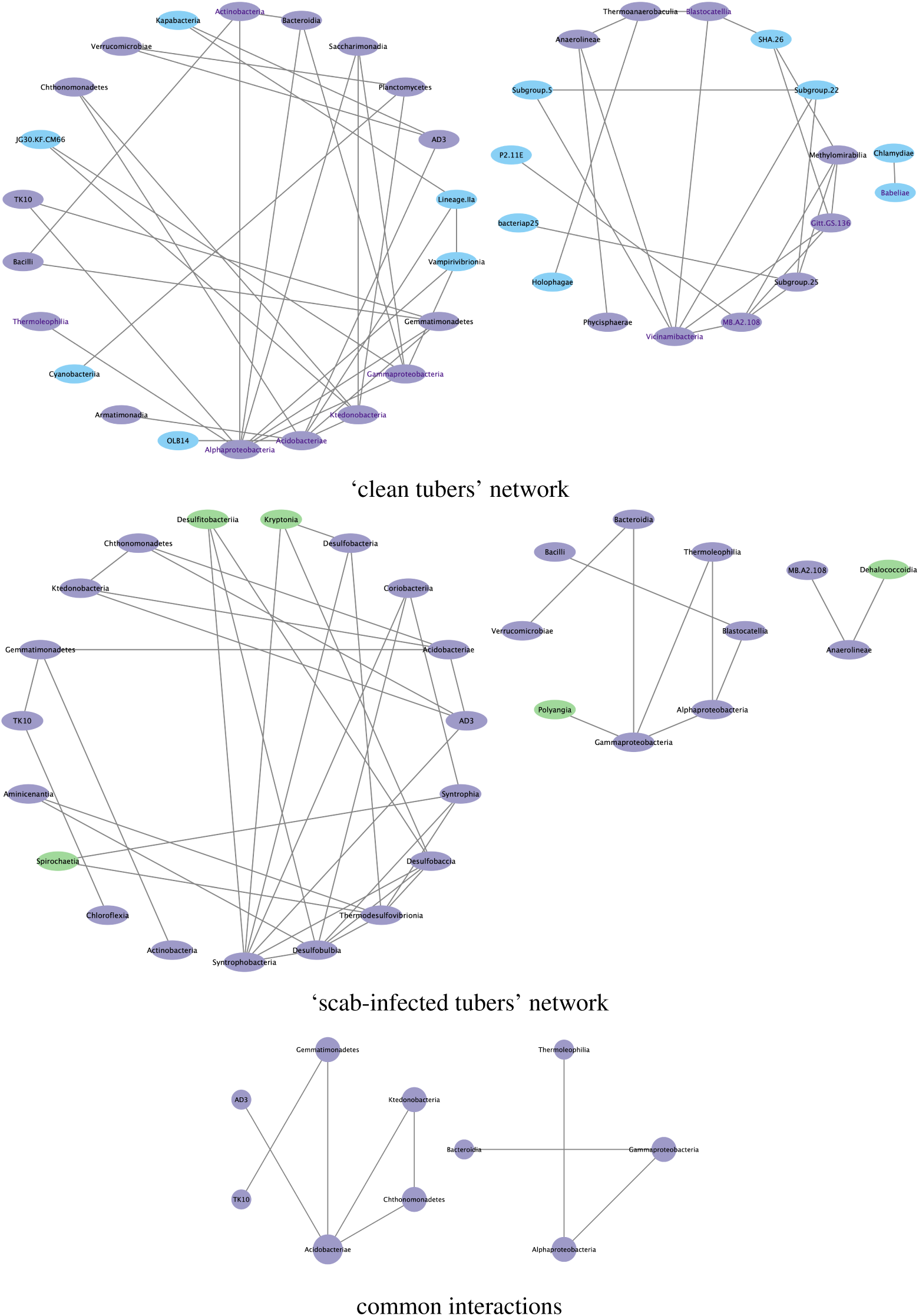
This figure illustrates the microbiome network at the Class taxonomic level. Part (a) represents the ‘clean tubers’ network, part (b) displays the ‘scab-infected tubers’ network, and part (c) shows the common interactions between them. Nodes correspond to Operational Taxonomic Units (OTUs) and are color-coded: purple for OTUs shared between ‘clean tubers’ and ‘scab-infected tubers’ networks, blue for OTUs unique to the ‘clean tubers’ network, and green for OTUs unique to the ‘scab-infected tubers’ network. Edges are shown by solid lines which confirmed by all four method).

**Figure S3:**
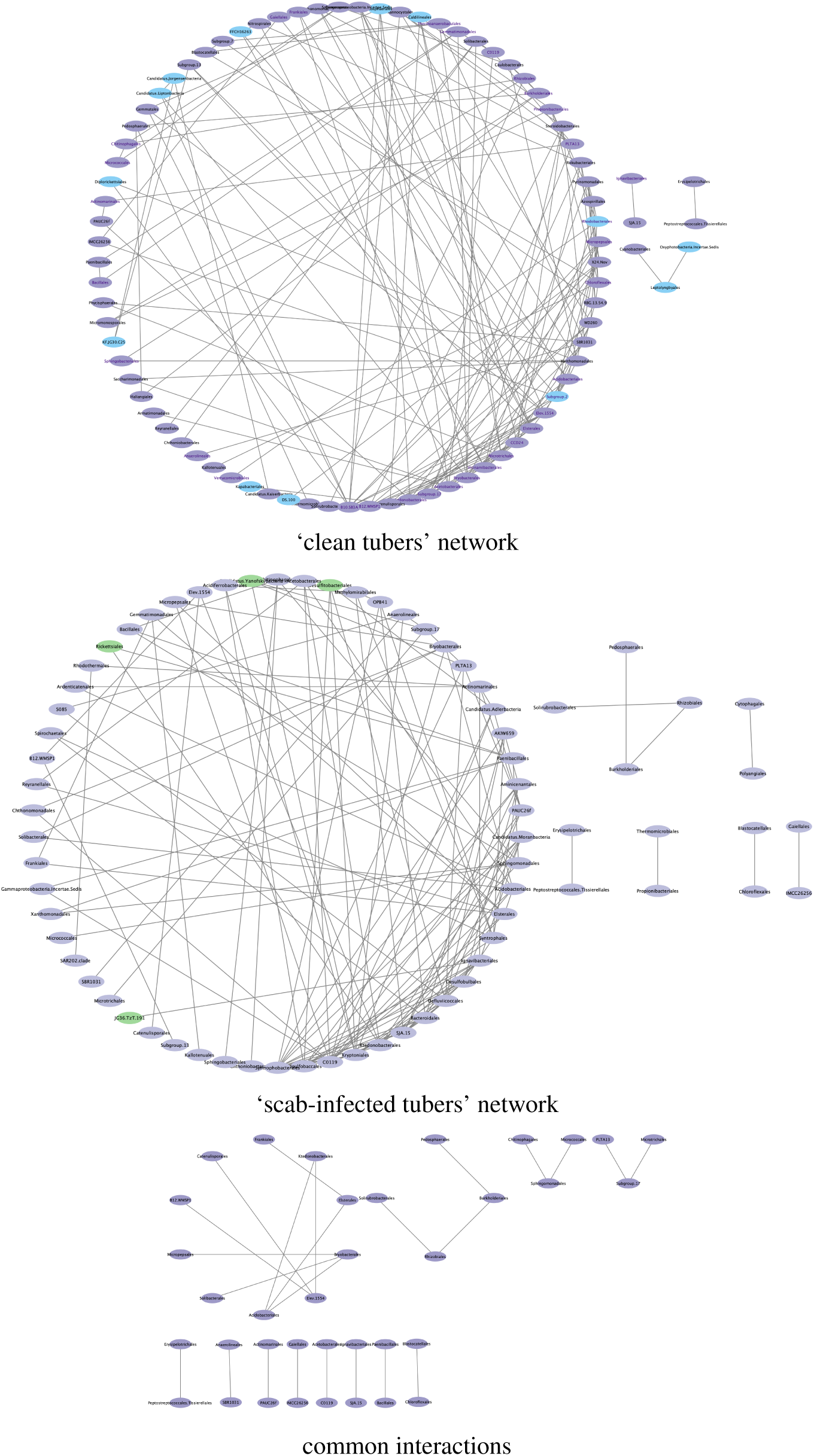
This figure showcases the microbiome network at the Order taxonomic level. Part (a) represents the ‘clean tubers’ network, part (b) displays the ‘scab-infected tubers’ network, and part (c) shows the common interactions between them. Nodes correspond to Operational Taxonomic Units (OTUs) and are color-coded: purple for OTUs shared between ‘clean tubers’ and ‘scab-infected tubers’ networks, blue for OTUs unique to the ‘clean tubers’ network, and green for OTUs unique to the ‘scab-infected tubers’ network. The top 20% of nodes with the highest centrality scores resulted by Equation (5) are labeled in dark purple. Node size reflects their degree of connectivity. Edges are shown as solid lines that confirmed by all four methods.

**Table 2:** Network metrics for microbiome networks constructed using four different methods based on all samples at the Phylum level.

| Method | num_edges | average_path_length | transitivity | mean_degree | mean_distance | modularity |
| --- | --- | --- | --- | --- | --- | --- |
| SPARCC | 148 | 1.917 | 0.506 | 3.895 | 1.917 | 0.238 |
| SE_glasso | 60 | 2.322 | 0.429 | 1.579 | 2.322 | 0.351 |
| SPRING | 172 | 2.658 | 0.272 | 4.526 | 2.658 | 0.456 |
| CMIMN | 136 | 3.06 | 0.636 | 3.579 | 3.06 | 0.372 |

Table 2 summarizes network metrics for different methods applied at the Phylum level of microbiome analysis. Notable variations in network characteristics are observed across these methods. SPRING constructs a network with 172 edges, indicating a relatively complex network structure, while SE_glasso results in a sparser network with only 60 edges. SPARCC falls in between with 148 edges, and CMIMN generates a network with 136 edges. In terms of average path length, SPARCC exhibits shorter paths on average (1.917), suggesting high connectivity, whereas CMIMN has longer average path lengths indicating less direct connections. Transitivity, a measure of clustering, is highest for CMIMN (0.636), implying a higher level of clustering in its network. The mean_degree and mean_distance offer insights into network density and structure; SPARCC and SPRING have higher mean degrees (3.895 and 4.526, respectively) and shorter mean distances (1.917 and 2.322, respectively), suggesting denser and more compact networks. Finally, SPRING exhibits the highest modularity value (0.456), indicating a potential modular structure within its network. These metrics collectively illustrate that different methods yield networks with distinct structural characteristics, highlighting the importance of selecting the method that aligns with specific research objectives and hypotheses.

**Table 3:** Network metrics for microbiome networks constructed using four different methods based on all samples at the Class level.

| topology_name | num_edges | average_path_length | transitivity | mean_degree | mean_distance | modularity |
| --- | --- | --- | --- | --- | --- | --- |
| SPARCC | 880 | 2.249 | 0.597 | 8.889 | 2.249 | 0.195 |
| SE_glasso | 228 | 2.388 | 0.562 | 2.303 | 2.388 | 0.333 |
| SPRING | 732 | 2.697 | 0.208 | 7.394 | 2.697 | 0.409 |
| CMIMN | 856 | 2.402 | 0.603 | 8.646 | 2.402 | 0.467 |

Table 3 provides network metrics for various methods applied at the Class level of microbiome analysis. Notable differences in network characteristics are evident across these methods. SPARCC yields a network with 880 edges, indicating a relatively complex and highly connected structure. SE_glasso, in contrast, results in a sparser network with 228 edges, suggesting fewer connections between Class-level components. SPRING’s network falls in between with 732 edges, while CMIMN constructs a network with 856 edges. When examining the average path length, SPARCC exhibits shorter paths on average (2.249), implying higher overall connectivity. SE_glasso and SPRING have longer average path lengths (2.388 and 2.697, respectively), indicating relatively less direct connections. Transitivity, a measure of network clustering, is highest for CMIMN (0.603), signifying a substantial degree of clustering within its network. Mean degree and mean distance provide insights into the density and overall structure of the networks; SPARCC and CMIMN have higher mean degrees (8.889 and 8.646, respectively) and shorter mean distances (2.249 and 2.402, respectively), suggesting denser and more compact networks. Finally, in terms of modularity, CMIMN exhibits the highest value (0.467), indicating a potential modular structure within its network.

**Table 4:** Network metrics for microbiome networks constructed using four different methods based on all samples at the Order level.

| topology_name | num_edges | average_path_length | transitivity | mean_degree | mean_distance | modularity |
| --- | --- | --- | --- | --- | --- | --- |
| SPARCC | 3400 | 2.091 | 0.647 | 17.989 | 2.091 | 0.212 |
| SE_glasso | 2170 | 2.414 | 0.503 | 11.481 | 2.414 | 0.324 |
| SPRING | 2060 | 2.595 | 0.175 | 10.899 | 2.595 | 0.399 |
| CMIMN | 3574 | 2.391 | 0.626 | 18.91 | 2.391 | 0.426 |

Table 4 presents network metrics for different methods applied at the Order level of microbiome analysis. Notable variations in network characteristics are observed across these methods. SPARCC produces a network with 3,400 edges, indicating a complex and highly connected structure. SE_glasso results in a network with 2,170 edges, suggesting a somewhat sparser network compared to SPARCC. SPRING falls in between with 2,060 edges, while CMIMN constructs a network with 3,574 edges, representing a highly connected network. Examining average path length, SPARCC exhibits shorter paths on average (2.091), indicating a high degree of connectivity. SE_glasso and SPRING have longer average path lengths (2.414 and 2.595, respectively), implying less direct connections. Transitivity, a measure of network clustering, is highest for SPARCC (0.647), signifying a substantial level of clustering within its network. Mean degree and mean distance provide insights into network density and structure; SPARCC and CMIMN have higher mean degrees (17.989 and 18.91, respectively) and shorter mean distances (2.091 and 2.391, respectively), suggesting dense and compact networks. Finally, in terms of modularity, CMIMN exhibits the highest value (0.426), indicating a potential modular structure within its network. These metrics collectively demonstrate that different methods yield networks with distinct structural characteristics at the Order level, emphasizing the importance of method selection based on specific research objectives and hypotheses.

In Tables S2 to S4, which present network metrics for various taxonomic levels, we observed notable trends. Specifically, the methods yielded distinct results:

- For average_path_length, mean_distance, and modularity, the SPARCC method consistently produced the lowest values, indicating its efficiency in minimizing these metrics across all taxonomic levels.
- Conversely, in the case of transitivity, the SPRING algorithm consistently yielded the lowest values. This suggests that the SPRING algorithm is particularly effective in minimizing transitivity, irrespective of the taxonomic level under consideration.

**Table 5:** Commonly identified important OTUs based on topological features in microbiome networks constructed from all samples at the **Phylum** level. Microbiome networks were constructed using four different inference methods: `SE_glasso`, `SPRING`, `SPARCC`, and `CMIMN`. For each network, key topological features—Degree, Betweenness, Closeness, Eigenvector Centrality, and PageRank—were computed to assess the importance of individual OTUs. The top 20% of OTUs based on each centrality measure were selected as important. An OTU was included in this table if it consistently ranked in the top 20% across all four network inference methods based on at least one topological feature. For example, *Proteobacteria* was ranked in the top 20% of networks generated by all four methods based on Degree, Closeness, and PageRank.

| Important OTUs | Features |
| --- | --- |
| Proteobacteria | Degree, Closeness, Page Rank |
| Acidobacteriota | Degree, Closeness, Page Rank |
| WPS.2 | Betweenness, Closeness |
| NB1.j | Page Rank |

**Table 6:** Commonly identified important OTUs based on topological features in microbiome networks constructed from all samples at the **Class** level. Microbiome networks were constructed using four different inference methods: `SE_glasso`, `SPRING`, `SPARCC`, and `CMIMN`. For each network, key topological features—Degree, Betweenness, Closeness, Eigenvector Centrality, and PageRank—were computed to assess the importance of individual OTUs. The top 20% of OTUs based on each centrality measure were selected as important. An OTU was included in this table if it consistently ranked in the top 20% across all four network inference methods based on at least one topological feature. For example, *Acidobacteriae* was consistently ranked in the top 20% of networks generated by all four methods based on all topological features.

| OTUs | Topology Measures |
| --- | --- |
| Acidobacteriae | Degree, Betweenness, Closeness, Eigenvector Centrality, Page Rank |
| Alphaproteobacteria | Degree, Betweenness, Closeness, Page Rank |
| Anaerolineae | Degree, Betweenness, Closeness |
| Ignavibacteria | Degree, Eigenvector Centrality, Page Rank |
| Ktedonobacteria | Betweenness |
| Gammaproteobacteria | Betweenness, Page Rank |
| MB.A2.108 | Closeness |
| OM190 | Closeness |
| Syntrophobacteria | Eigenvector Centrality |
| Kryptonia | Eigenvector Centrality |
| Desulfobulbia | Eigenvector Centrality |

**Table 7:**
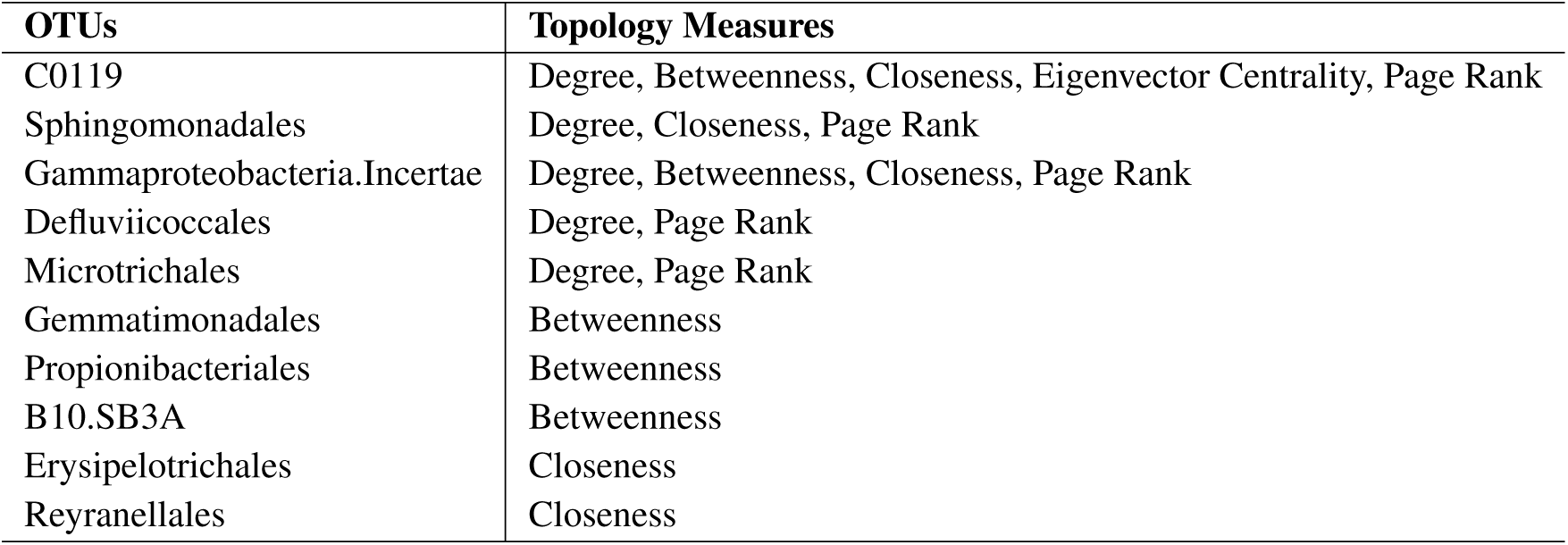
Commonly identified important OTUs based on topological features in microbiome networks constructed from all samples at the **Order** level. Microbiome networks were constructed using four different inference methods: `SE_glasso`, `SPRING`, `SPARCC`, and `CMIMN`. For each network, key topological features—Degree, Betweenness, Closeness, Eigenvector Centrality, and PageRank—were computed to assess the importance of individual OTUs. The top 20% of OTUs based on each centrality measure were selected as important. An OTU was included in this table if it consistently ranked in the top 20% across all four network inference methods based on at least one topological feature. For example, *C0119* was consistently ranked in the top 20% of networks generated by all four methods based on all topological features.

**Table 8:**
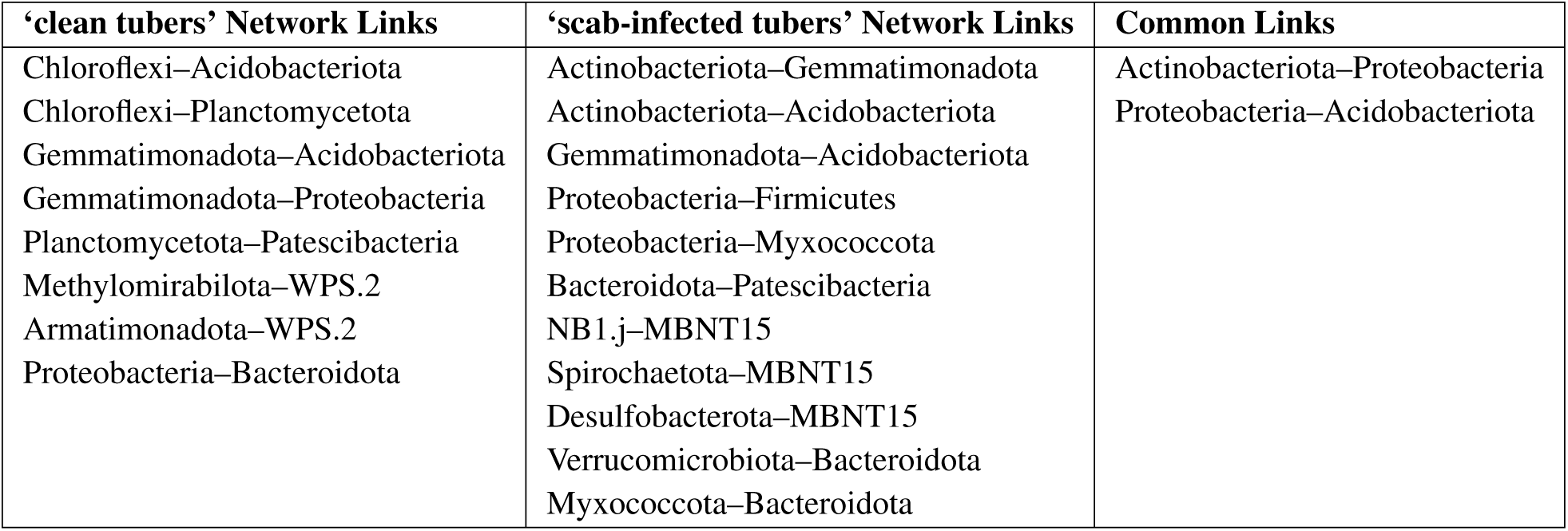
Phylum-level microbial associations identified in ‘clean tuber’ and ‘scab-infected tubers’ microbiome networks. Only interactions confirmed by all four inference methods are shown.

| ‘clean tubers’ Network Links | ‘scab-infected tubers’ Network Links | Common Links |
| --- | --- | --- |
| Chloroflexi–Acidobacteriota<br>Chloroflexi–Planctomycetota<br>Gemmatimonadota–Acidobacteriota<br>Gemmatimonadota–Proteobacteria<br>Planctomycetota–Patescibacteria<br>Methyloirabilota–WPS.2<br>Armatimonadota–WPS.2<br>Proteobacteria–Bacteroidota | Actinobacteriota–Gemmatimonadota<br>Actinobacteriota–Acidobacteriota<br>Gemmatimonadota–Acidobacteriota<br>Proteobacteria–Firmicutes<br>Proteobacteria–Myxococcota<br>Bacteroidota–Patescibacteria<br>NB1.j–MBNT15<br>Spirochaetota–MBNT15<br>Desulfobacterota–MBNT15<br>Verrucomicrobiota–Bacteroidota<br>Myxococcota–Bacteroidota | Actinobacteriota–Proteobacteria<br>Proteobacteria–Acidobacteriota |

**Table 9:** Class-level microbial associations identified in ‘clean tuber’ and ‘scab-infected tubers’ microbiome networks. Only interactions confirmed by all four inference methods are shown.

| ‘clean tuber’ Network Links | ‘scab-infected tubers’ Network Links | Common Links |
| --- | --- | --- |
| AD3–Kapabacteria<br>AD3–Verrucomicrobiae<br>Acidobacteriae–Armatimonadia<br>Acidobacteriae–Lineage.IIa<br>Acidobacteriae–OLB14<br>Actinobacteria–Alphaproteobacteria<br>Actinobacteria–Bacilli<br>Actinobacteria–Bacteroidia<br>Alphaproteobacteria–Bacteroidia<br>Alphaproteobacteria–Gemmatimonadetes<br>Alphaproteobacteria–Saccharimonadia<br>Alphaproteobacteria–TK10<br>Alphaproteobacteria–Vampirivibrionia<br>Anaerolineae–Phycisphaerae<br>Anaerolineae–Thermoanaerobaculia<br>Anaerolineae–Vicinamibacteria<br>Babeliae–Chlamydiae<br>Bacilli–Gemmatimonadetes<br>Blastocatellia–SHA.26<br>Blastocatellia–Thermoanaerobaculia<br>Blastocatellia–Vicinamibacteria<br>Cyanobacteriia–Planctomycetes<br>Gammaproteobacteria–JG30.KF.CM66<br>Gammaproteobacteria–Saccharimonadia<br>Gammaproteobacteria–Vampirivibrionia<br>Gitt.GS.136–MB.A2.108<br>Gitt.GS.136–Methyloirabilia<br>Gitt.GS.136–SHA.26<br>Gitt.GS.136–Vicinamibacteria<br>Holophagae–Thermoanaerobaculia<br>JG30.KF.CM66–Ktedonobacteria<br>Kapabacteria–Lineage.IIa<br>Ktedonobacteria–Planctomycetes<br>Ktedonobacteria–Saccharimonadia<br>Lineage.IIa–Vampirivibrionia<br>MB.A2.108–Methyloirabilia<br>MB.A2.108–P2.11E<br>MB.A2.108–Subgroup.25<br>MB.A2.108–Vicinamibacteria<br>Methyloirabilia–SHA.26<br>Methyloirabilia–Subgroup.25 | AD3–Chthonomonadetes<br>AD3–Ktedonobacteria<br>AD3–Syntrophobacteria<br>Actinobacteria–Gemmatimonadetes<br>Alphaproteobacteria–Blastocatellia<br>Aminicenantia–Desulfobulbia<br>Aminicenantia–Thermodesulfovibrionia<br>Anaerolineae–Dehalococcoidia<br>Anaerolineae–MB.A2.108<br>Bacilli–Blastocatellia<br>Bacteroidia–Verrucomicrobiae<br>Chloroflexia–TK10<br>Coriobacteriia–Desulfobulbia<br>Coriobacteriia–Syntrophia<br>Coriobacteriia–Syntrophobacteria<br>Desulfitobacteriia–Desulfobaccia<br>Desulfitobacteriia–Desulfobulbia<br>Desulfitobacteriia–Syntrophobacteria<br>Desulfobaccia–Desulfobulbia<br>Desulfobaccia–Kryptonia<br>Desulfobaccia–Syntrophobacteria<br>Desulfobaccia–Thermodesulfovibrionia<br>Desulfobacteria–Kryptonia<br>Desulfobacteria–Syntrophobacteria<br>Desulfobacteria–Thermodesulfovibrionia<br>Desulfobulbia–Syntrophia<br>Desulfobulbia–Syntrophobacteria<br>Desulfobulbia–Thermodesulfovibrionia<br>Gammaproteobacteria–Polyangia<br>Gammaproteobacteria–Thermoleophilia<br>Kryptonia–Syntrophobacteria<br>Spirochaetia–Syntrophia<br>Spirochaetia–Thermodesulfovibrionia<br>Syntrophia–Thermodesulfovibrionia | AD3–Acidobacteriae<br>Acidobacteriae–Chthonomonadetes<br>Acidobacteriae–Gemmatimonadetes<br>Acidobacteriae–Ktedonobacteria<br>Alphaproteobacteria–Gammaproteobacteria<br>Alphaproteobacteria–Thermoleophilia<br>Bacteroidia–Gammaproteobacteria<br>Chthonomonadetes–Ktedonobacteria<br>Gemmatimonadetes–TK10 |

| Clean Network Links | Disease Network Links | Common Links |
| --- | --- | --- |
| Planctomycetes–Verrucomicrobiae<br>Subgroup.22–Subgroup.25<br>Subgroup.22–Subgroup.5<br>Subgroup.22–Vicinamibacteria<br>Subgroup.25–bacteriap25<br>Subgroup.5–Vicinamibacteria |  |  |

**Table 10:**
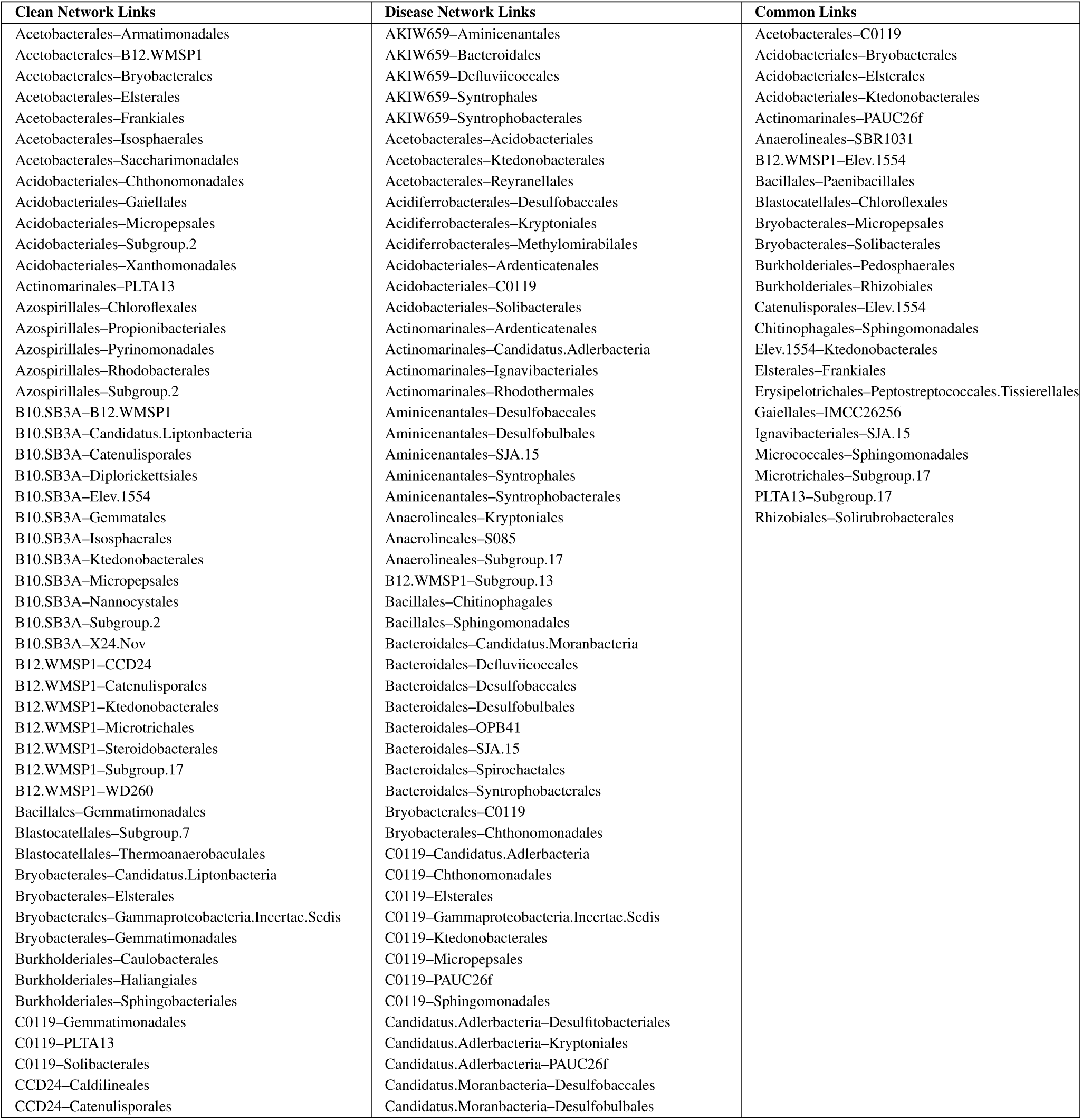
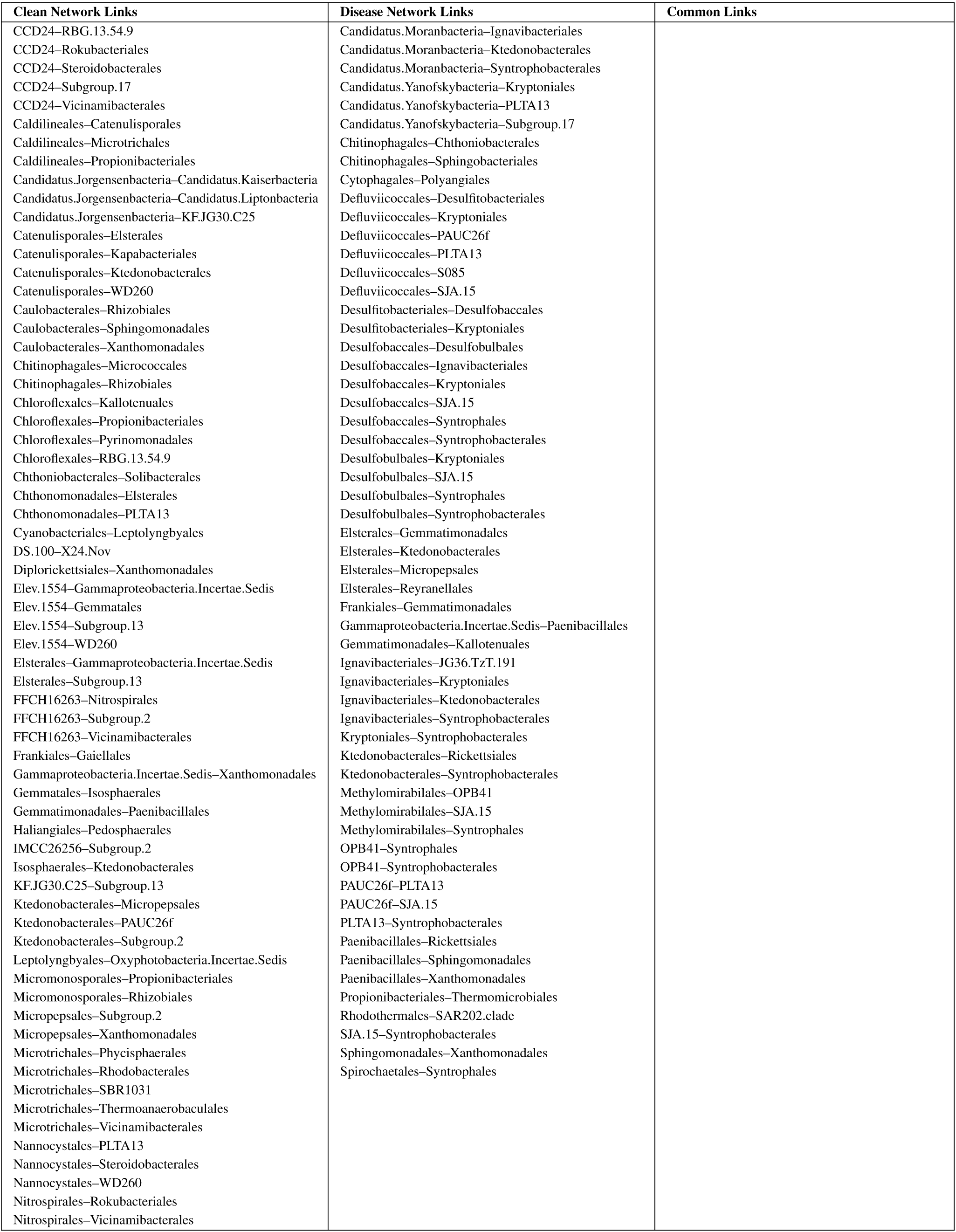
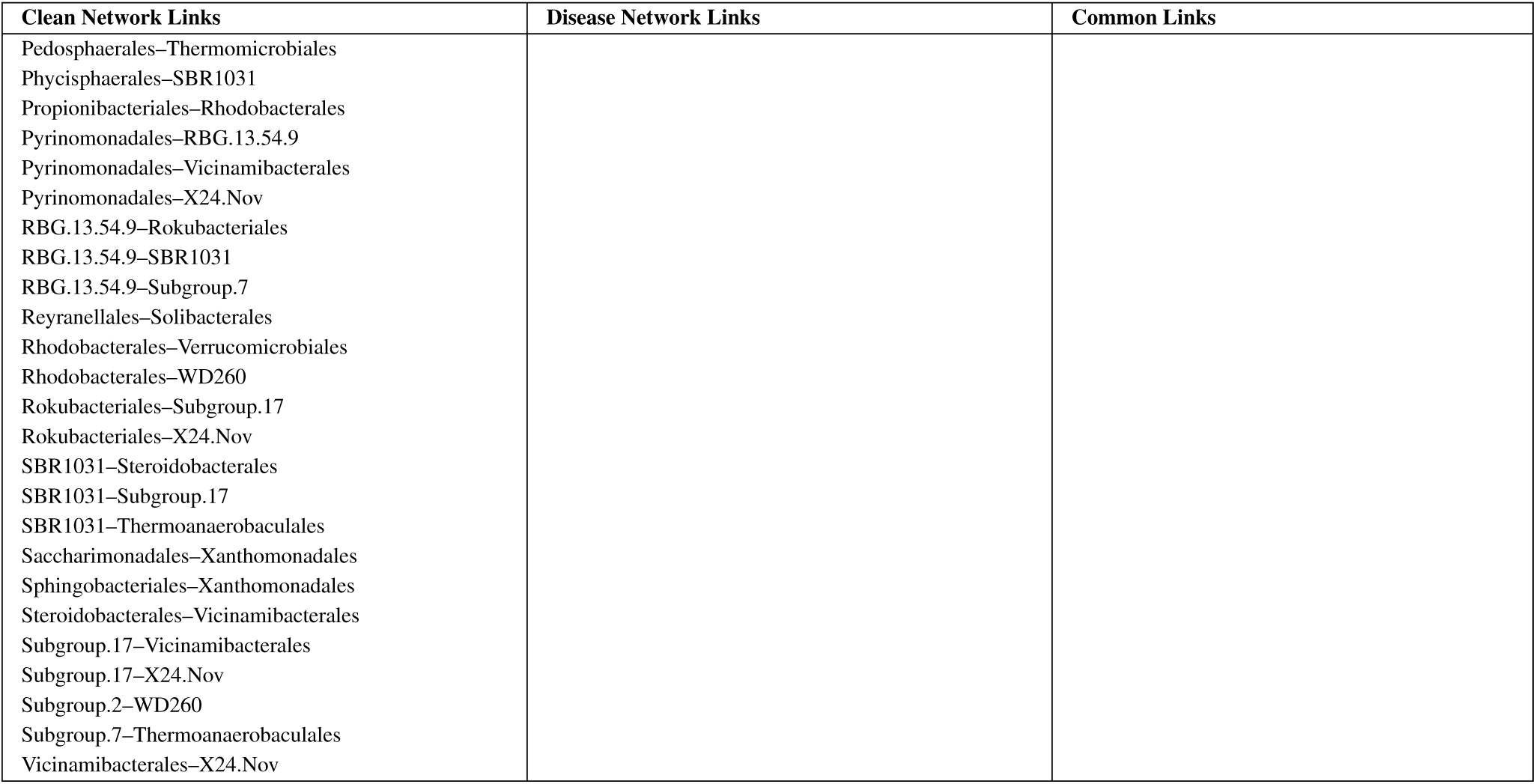
Order-level microbial associations identified in ‘clean tuber’ and ‘scab-infected tubers’ microbiome networks. Only interactions confirmed by all four inference methods are shown.

| Clean Network Links | Disease Network Links | Common Links |
| --- | --- | --- |
| Acetobacterales–Armatimonadales<br>Acetobacterales–B12.WMSP1<br>Acetobacterales–Bryobacterales<br>Acetobacterales–Elsterales<br>Acetobacterales–Frankiales<br>Acetobacterales–Isosphaerales<br>Acetobacterales–Saccharimonadales<br>Acidobacteriales–Chthonomonadales<br>Acidobacteriales–Gaiellales<br>Acidobacteriales–Micropepsales<br>Acidobacteriales–Subgroup.2<br>Acidobacteriales–Xanthomonadales<br>Actinomarinales–PLTA13<br>Azospirillales–Chloroflexales<br>Azospirillales–Propionibacteriales<br>Azospirillales–Pyrenomonadales<br>Azospirillales–Rhodobacterales<br>Azospirillales–Subgroup.2<br>B10.SB3A–B12.WMSP1<br>B10.SB3A–Candidatus.Liptonbacteria<br>B10.SB3A–Catenulisporales<br>B10.SB3A–Diploricetisiales<br>B10.SB3A–Elev.1554<br>B10.SB3A–Gemmatales<br>B10.SB3A–Isosphaerales<br>B10.SB3A–Ktedonobacterales<br>B10.SB3A–Micropepsales<br>B10.SB3A–Nannocystales<br>B10.SB3A–Subgroup.2<br>B10.SB3A–X24.Nov<br>B12.WMSP1–CCD24<br>B12.WMSP1–Catenulisporales<br>B12.WMSP1–Ktedonobacterales<br>B12.WMSP1–Microtrichales<br>B12.WMSP1–Steroidobacterales<br>B12.WMSP1–Subgroup.17<br>B12.WMSP1–WD260<br>Bacillales–Gemmatimonadales<br>Blastocatellales–Subgroup.7<br>Blastocatellales–Thermoanaerobaculales<br>Bryobacteriales–Candidatus.Liptonbacteria<br>Bryobacteriales–Elsterales<br>Bryobacteriales–Gammaproteobacteria.Incertae.Sedis<br>Bryobacteriales–Gemmatimonadales<br>Burkholderiales–Caulobacteriales<br>Burkholderiales–Haliangiales<br>Burkholderiales–Sphingobacteriales<br>C0119–Gemmatimonadales<br>C0119–PLTA13<br>C0119–Solibacteriales<br>CCD24–Caldilineales<br>CCD24–Catenulisporales | AKIW659–Aminicenantales<br>AKIW659–Bacteroidales<br>AKIW659–Deffluviococcales<br>AKIW659–Syntrophales<br>AKIW659–Syntrophobacteriales<br>Acetobacterales–Acidobacteriales<br>Acetobacterales–Ktedonobacteriales<br>Acetobacterales–Reyranellales<br>Acidiferrobacteriales–Desulfobaccales<br>Acidiferrobacteriales–Kryptoniales<br>Acidiferrobacteriales–Methylomirabilales<br>Acidobacteriales–Ardenticatenales<br>Acidobacteriales–C0119<br>Acidobacteriales–Solibacteriales<br>Actinomarinales–Ardenticatenales<br>Actinomarinales–Candidatus.Adlerbacteria<br>Actinomarinales–Ignavibacteriales<br>Actinomarinales–Rhodothermales<br>Aminicenantales–Desulfobaccales<br>Aminicenantales–Desulfobulbales<br>Aminicenantales–SJA.15<br>Aminicenantales–Syntrophales<br>Aminicenantales–Syntrophobacteriales<br>Anaerolineales–Kryptoniales<br>Anaerolineales–S085<br>Anaerolineales–Subgroup.17<br>B12.WMSP1–Subgroup.13<br>Bacillales–Chitinophagales<br>Bacillales–Sphingomonadales<br>Bacteroidales–Candidatus.Moranbacteria<br>Bacteroidales–Deffluviococcales<br>Bacteroidales–Desulfobaccales<br>Bacteroidales–Desulfobulbales<br>Bacteroidales–OPB41<br>Bacteroidales–SJA.15<br>Bacteroidales–Spirochaetales<br>Bacteroidales–Syntrophobacteriales<br>Bryobacteriales–C0119<br>Bryobacteriales–Chthonomonadales<br>C0119–Candidatus.Adlerbacteria<br>C0119–Chthonomonadales<br>C0119–Elsterales<br>C0119–Gammaproteobacteria.Incertae.Sedis<br>C0119–Ktedonobacteriales<br>C0119–Micropepsales<br>C0119–PAUC26f<br>C0119–Sphingomonadales<br>Candidatus.Adlerbacteria–Desulfobacteriales<br>Candidatus.Adlerbacteria–Kryptoniales<br>Candidatus.Adlerbacteria–PAUC26f<br>Candidatus.Moranbacteria–Desulfobaccales<br>Candidatus.Moranbacteria–Desulfobulbales | Acetobacterales–C0119<br>Acidobacteriales–Bryobacteriales<br>Acidobacteriales–Elsterales<br>Acidobacteriales–Ktedonobacteriales<br>Actinomarinales–PAUC26f<br>Anaerolineales–SBR1031<br>B12.WMSP1–Elev.1554<br>Bacillales–Paenibacillales<br>Blastocatellales–Chloroflexales<br>Bryobacteriales–Micropepsales<br>Bryobacteriales–Solibacteriales<br>Burkholderiales–Pedosphaerales<br>Burkholderiales–Rhizobiales<br>Catenulisporales–Elev.1554<br>Chitinophagales–Sphingomonadales<br>Elev.1554–Ktedonobacteriales<br>Elsterales–Frankiales<br>Erysipelotrichales–Peptostreptococcales.Tissierellales<br>Gaiellales–IMCC26256<br>Ignavibacteriales–SJA.15<br>Micrococcales–Sphingomonadales<br>Microtrichales–Subgroup.17<br>PLTA13–Subgroup.17<br>Rhizobiales–Solirubrobacteriales |
| CCD24–RBG.13.54.9<br>CCD24–Rokubacteriales<br>CCD24–Steroidobacteriales<br>CCD24–Subgroup.17<br>CCD24–Vicinamibacteriales<br>Caldilineales–Catenulisporales<br>Caldilineales–Microtrichales<br>Caldilineales–Propionibacteriales<br>Candidatus.Jorgensenbacteria–Candidatus.Kaiserbacteria<br>Candidatus.Jorgensenbacteria–Candidatus.Liptonbacteria<br>Candidatus.Jorgensenbacteria–KF.JG30.C25<br>Catenulisporales–Elsterales<br>Catenulisporales–Kapabacteriales<br>Catenulisporales–Ktedonobacteriales<br>Catenulisporales–WD260<br>Caulobacteriales–Rhizobiales<br>Caulobacteriales–Sphingomonadales<br>Caulobacteriales–Xanthomonadales<br>Chitinophagales–Micrococcales<br>Chitinophagales–Rhizobiales<br>Chloroflexales–Kallotenuales<br>Chloroflexales–Propionibacteriales<br>Chloroflexales–Pyrimonadales<br>Chloroflexales–RBG.13.54.9<br>Chthoniobacteriales–Solibacteriales<br>Chthonomonadales–Elsterales<br>Chthonomonadales–PLTA13<br>Cyanobacteriales–Leptolyngbyales<br>DS.100–X24.Nov<br>Diplorickettsiales–Xanthomonadales<br>Elev.1554–Gammaproteobacteria.Incertae.Sedis<br>Elev.1554–Gemmatales<br>Elev.1554–Subgroup.13<br>Elev.1554–WD260<br>Elsterales–Gammaproteobacteria.Incertae.Sedis<br>Elsterales–Subgroup.13<br>FFCH16263–Nitrospirales<br>FFCH16263–Subgroup.2<br>FFCH16263–Vicinamibacteriales<br>Frankiales–Gaiellales<br>Gammaproteobacteria.Incertae.Sedis–Xanthomonadales<br>Gemmatales–Isosphaerales<br>Gemmatimonadales–Paenibacillales<br>Haliangiales–Pedosphaerales<br>IMCC26256–Subgroup.2<br>Isosphaerales–Ktedonobacteriales<br>KF.JG30.C25–Subgroup.13<br>Ktedonobacteriales–Micropepsales<br>Ktedonobacteriales–PAUC26f<br>Ktedonobacteriales–Subgroup.2<br>Leptolyngbyales–Oxyphotobacteria.Incertae.Sedis<br>Micromonosporales–Propionibacteriales<br>Micromonosporales–Rhizobiales<br>Micropepsales–Subgroup.2<br>Micropepsales–Xanthomonadales<br>Microtrichales–Phycisphaerales<br>Microtrichales–Rhodobacteriales<br>Microtrichales–SBR1031<br>Microtrichales–Thermoanaerobaculales<br>Microtrichales–Vicinamibacteriales<br>Nannocystales–PLTA13<br>Nannocystales–Steroidobacteriales<br>Nannocystales–WD260<br>Nitrospirales–Rokubacteriales<br>Nitrospirales–Vicinamibacteriales | Candidatus.Moranbacteria–Ignavibacteriales<br>Candidatus.Moranbacteria–Ktedonobacteriales<br>Candidatus.Moranbacteria–Syntrophobacteriales<br>Candidatus.Yanofskybacteria–Kryptoniales<br>Candidatus.Yanofskybacteria–PLTA13<br>Candidatus.Yanofskybacteria–Subgroup.17<br>Chitinophagales–Chthoniobacteriales<br>Chitinophagales–Sphingobacteriales<br>Cytophagales–Polyangiales<br>Defluviococcales–Desulfotobacteriales<br>Defluviococcales–Kryptoniales<br>Defluviococcales–PAUC26f<br>Defluviococcales–PLTA13<br>Defluviococcales–S085<br>Defluviococcales–SJA.15<br>Desulfotobacteriales–Desulfobaccales<br>Desulfotobacteriales–Kryptoniales<br>Desulfobaccales–Desulfobulbales<br>Desulfobaccales–Ignavibacteriales<br>Desulfobaccales–Kryptoniales<br>Desulfobaccales–SJA.15<br>Desulfobaccales–Syntrophales<br>Desulfobaccales–Syntrophobacteriales<br>Desulfobulbales–Kryptoniales<br>Desulfobulbales–SJA.15<br>Desulfobulbales–Syntrophales<br>Desulfobulbales–Syntrophobacteriales<br>Elsterales–Gemmatimonadales<br>Elsterales–Ktedonobacteriales<br>Elsterales–Micropepsales<br>Elsterales–Reyrnellales<br>Frankiales–Gemmatimonadales<br>Gammaproteobacteria.Incertae.Sedis–Paenibacillales<br>Gemmatimonadales–Kallotenuales<br>Ignavibacteriales–JG36.TzT.191<br>Ignavibacteriales–Kryptoniales<br>Ignavibacteriales–Ktedonobacteriales<br>Ignavibacteriales–Syntrophobacteriales<br>Kryptoniales–Syntrophobacteriales<br>Ktedonobacteriales–Rickettsiales<br>Ktedonobacteriales–Syntrophobacteriales<br>Methylomirabilales–OPB41<br>Methylomirabilales–SJA.15<br>Methylomirabilales–Syntrophales<br>OPB41–Syntrophales<br>OPB41–Syntrophobacteriales<br>PAUC26f–PLTA13<br>PAUC26f–SJA.15<br>PLTA13–Syntrophobacteriales<br>Paenibacillales–Rickettsiales<br>Paenibacillales–Sphingomonadales<br>Paenibacillales–Xanthomonadales<br>Propionibacteriales–Thermomicrobiales<br>Rhodothermales–SAR202.clade<br>SJA.15–Syntrophobacteriales<br>Sphingomonadales–Xanthomonadales<br>Spirochaetales–Syntrophales |  |
| Pedosphaerales–Thermomicrobiales<br>Phycisphaerales–SBR1031<br>Propionibacteriales–Rhodobacterales<br>Pyrinomonadales–RBG.13.54.9<br>Pyrinomonadales–Vicinamibacteriales<br>Pyrinomonadales–X24.Nov<br>RBG.13.54.9–Rokubacteriales<br>RBG.13.54.9–SBR1031<br>RBG.13.54.9–Subgroup.7<br>Reyranelles–Solibacteriales<br>Rhodobacterales–Verrucomicrobiales<br>Rhodobacterales–WD260<br>Rokubacteriales–Subgroup.17<br>Rokubacteriales–X24.Nov<br>SBR1031–Steroidobacteriales<br>SBR1031–Subgroup.17<br>SBR1031–Thermoanaerobaculales<br>Saccharimonadales–Xanthomonadales<br>Sphingobacteriales–Xanthomonadales<br>Steroidobacteriales–Vicinamibacteriales<br>Subgroup.17–Vicinamibacteriales<br>Subgroup.17–X24.Nov<br>Subgroup.2–WD260<br>Subgroup.7–Thermoanaerobaculales<br>Vicinamibacteriales–X24.Nov |  |  |

**Table 11:** Important OTUs identified using Multi Machine Learning (ML) methods at the **Phylum** level. Microbiome data were normalized using four different techniques: (1) Centered Log-Ratio (CLR), (2) the original dataset (no transformation), (3) Log transformation, and (4) Total Sum Scaling (TSS). Seven ML-based feature selection methods were applied to each normalized dataset to identify important OTUs. An OTU was selected if it was identified as important by at least five out of the seven ML methods. The table presents the OTUs identified as important for each normalization method, with separate columns for ML_Clr (CLR-normalized data), ML_Original (original dataset), ML_Log (log-transformed data), and ML_TSS (TSS-normalized data). The final ML_Intersection column lists the OTUs that were consistently selected as important across all normalization methods, highlighting robust microbial taxa that are independent of the normalization approach.

| ML_Clr | ML_Original | ML_Log | ML_TSS | ML_Intersection |
| --- | --- | --- | --- | --- |
| Firmicutes | Armatimonadota | Firmicutes | Firmicutes | Firmicutes |
| Cyanobacteria | Firmicutes | Armatimonadota | Cyanobacteria | Cyanobacteria |
| Methylomirabilota | Deinococcota | Cyanobacteria | Patescibacteria | Armatimonadota |
| Armatimonadota | Cyanobacteria | Deinococcota | Armatimonadota | NB1.j |
| Deinococcota | NB1.j | Verrucomicrobiota | Bacteroidota |  |
| WPS.2 | Verrucomicrobiota | Spirochaetota | Spirochaetota |  |
| Acidobacteriota | Patescibacteria | NB1.j | Methylomirabilota |  |
| MBNT15 | Methylomirabilota | Bacteroidota | Acidobacteriota |  |
| NB1.j |  | WPS.2 | Nitrospirota |  |
| Bacteroidota |  |  | Desulfobacterota |  |
| Verrucomicrobiota |  |  | NB1.j |  |

**Table 12:** Important OTUs identified using Multi Machine Learning (ML) methods at the **Class** level. Microbiome data were normalized using four different techniques: (1) Centered Log-Ratio (CLR), (2) the original dataset (no transformation), (3) Log transformation, and (4) Total Sum Scaling (TSS). Seven ML-based feature selection methods were applied to each normalized dataset to identify important OTUs. An OTU was selected if it was identified as important by at least five out of the seven ML methods. The table presents the OTUs identified as important for each normalization method, with separate columns for ML_Clr (CLR-normalized data), ML_Original (original dataset), ML_Log (log-transformed data), and ML_TSS (TSS-normalized data). The final ML_Intersection column lists the OTUs that were consistently selected as important across all normalization methods, highlighting robust microbial taxa that are independent of the normalization approach.

| ML_Clr | ML_Original | ML_Log | ML_TSS | ML_Intersection |
| --- | --- | --- | --- | --- |
| Bacilli | Bacilli | Bacilli | Vicinamibacteria | Bacilli |
| Ktedonobacteria | Latescibacteria | Ignavibacteria | Saccharimonadia | Ktedonobacteria |
| Cyanobacteriia | Saccharimonadia | Verrucomicrobiae | Gemmatimonadetes | Cyanobacteriia |
| Saccharimonadia | Ktedonobacteria | Deinococci | Gitt.GS.136 | Saccharimonadia |
| Planctomycetes | Spirochaetia | Latescibacteria | Oligoflexia | Planctomycetes |
| Lineage.IIa | Planctomycetes | Saccharimonadia | Ktedonobacteria | Ignavibacteria |
| Vampirivibrionia | Kryptonia | Ktedonobacteria | Anaerolineae | Dehalococcoidia |
| Verrucomicrobiae | Verrucomicrobiae | OLB14 | Bacilli | Anaerolineae |
| Ignavibacteria | Deinococci | Kryptonia | Acidimicrobiia | MB.A2.108 |
| Dehalococcoidia | Ignavibacteria | MB.A2.108 | Planctomycetes | Chthonomonadetes |
| Thermoplasmata | OLB14 | Cyanobacteriia | Nitrospiria | Kryptonia |
| Deinococci | Cyanobacteriia | Planctomycetes | Methylomirabilia |  |
| Anaerolineae | Dehalococcoidia | AD3 | Ignavibacteria |  |
| Pla4.lineage | Methylomirabilia | Gitt.GS.136 | MB.A2.108 |  |
| MB.A2.108 | Chthonomonadetes | Dehalococcoidia | Acidobacteriae |  |
| Methylomirabilia | MB.A2.108 | Anaerolineae | TK10 |  |
| Chthonomonadetes | Gitt.GS.136 | Armatimonadia | Bacteroidia |  |
| OLB14 | Vicinamibacteria | Chthonomonadetes | Polyangia |  |
| Bacteroidia | Anaerolineae | Oligoflexia | Thermoleophilia |  |
| Oligoflexia | Vampirivibrionia | Clostridia | KD4.96 |  |
| Kryptonia | Holophagae | Spirochaetia | Cyanobacteriia |  |
| TK10 |  |  | Chthonomonadetes |  |
|  |  |  | Dehalococcoidia |  |
|  |  |  | Kryptonia |  |

**Table 13:** Important OTUs identified using Multi Machine Learning (ML) methods at the **Order** level. Microbiome data were normalized using four different techniques: (1) Centered Log-Ratio (CLR), (2) the original dataset (no transformation), (3) Log transformation, and (4) Total Sum Scaling (TSS). Seven ML-based feature selection methods were applied to each normalized dataset to identify important OTUs. An OTU was selected if it was identified as important by at least five out of the seven ML methods. The table presents the OTUs identified as important for each normalization method, with separate columns for ML_Clr (CLR-normalized data), ML_Original (original dataset), ML_Log (log-transformed data), and ML_TSS (TSS-normalized data). The final ML_Intersection column lists the OTUs that were consistently selected as important across all normalization methods, highlighting robust microbial taxa that are independent of the normalization approach.

| ML_Clr | ML_Original | ML_Log | ML_TSS | ML_Intersection |
| --- | --- | --- | --- | --- |
| Saccharimonadales | Anaerolineales | Saccharimonadales | Sphingomonadales | Saccharimonadales |
| Bacillales | Acetobacterales | Bacillales | Saccharimonadales | Bacillales |
| C0119 | Chloroplast | Subgroup.2 | SBR1031 | C0119 |
| Isosphaerales | PAUC26f | Solibacteriales | Kryptoniales | Subgroup.2 |
| Subgroup.2 | Subgroup.17 | Micropepsales | Vicinamibacteriales | Xanthomonadales |
| Blastocatellales | Bacteroidales | Anaerolineales | Anaerolineales | Acidobacteriales |
| Xanthomonadales | Alicyclobacillales | Latescibacteriales | C0119 | Chloroplast |
| Acidobacteriales | Micropepsales | Frankiales | Bryobacteriales | Alicyclobacillales |
| Chloroplast | Bacillales | Gaiellales | Micropepsales | Paenibacillales |
| Alicyclobacillales | Sphingomonadales | Alicyclobacillales | Chloroplast | Acetobacteriales |
| Paenibacillales | Defluviicoccales | Bacteroidales | Subgroup.2 | Pseudomonadales |
| Acetobacterales | Frankiales | B10.SB3A | Gaiellales | Anaerolineales |
| Pseudomonadales | Saccharimonadales | Acetobacterales | PLTA13 | Elsterales |
| Anaerolineales | Caulobacteriales | Chloroplast | Bacillales | Bacteroidales |
| Elsterales | Acidobacteriales | Defluviicoccales | Microtrichales | Ktedonobacteriales |
| Oligoflexales | Pseudomonadales | Propionibacteriales | Candidatus.Levyacteria | Sphingomonadales |
| Bacteroidales | Bryobacteriales | Acidobacteriales | Pseudomonadales | Kineosporiales |
| Deinococcales | C0119 | Sphingomonadales | Rokubacteriales | SBR1031 |
| Ktedonobacteriales | Gaiellales | Caulobacteriales | Acetobacterales | Rokubacteriales |
| Sphingomonadales | B10.SB3A | B12.WMSP1 | Clostridiales | Frankiales |
| Chloroflexales | Microtrichales | SBR1031 | PAUC26f | Micropepsales |
| Rhodobacteriales | Chthoniobacteriales | Chthoniobacteriales | Elsterales | Gaiellales |
| Candidatus.Yanofskybacteria | SBR1031 | Deinococcales | Frankiales | PLTA13 |
| KF.JG30.C25 | Kineosporiales | Pseudomonadales | B10.SB3A | Defluviicoccales |
| Kineosporiales | Ktedonobacteriales | Obscuribacteriales | Kineosporiales | Obscuribacteriales |
| SBR1031 | Subgroup.2 | Rokubacteriales | Ktedonobacteriales |  |
| Rokubacteriales | Latescibacteriales | Subgroup.17 | Subgroup.17 |  |
| Frankiales | Chitinophagales | Lineage.IV | Lineage.IV |  |
| Rickettsiales | B12.WMSP1 | PAUC26f | Micromonosporales |  |
| Micropepsales | Blastocatellales | Spirochaetales | Gammaproteobacteria.Incertae |  |
| Enterobacteriales | Xanthomonadales | Chitinophagales | Blastocatellales |  |
| Micrococcales | Syntrophobacteriales | Kineosporiales | Cytophagales |  |
| Gaiellales | Sphingobacteriales | Isosphaerales | Obscuribacteriales |  |
| PLTA13 | SJA.15 | Paenibacillales | Deinococcales |  |
| Solibacteriales | Elsterales | Xanthomonadales | Chthoniobacteriales |  |
| CCD24 | Pseudonocardiales | C0119 | Chthonomonadales |  |
| Defluviicoccales | Clostridiales | Ktedonobacteriales | Acidobacteriales |  |
| Desulfotobacteriales | Candidatus.Yanofskybacteria | Micromonosporales | Pseudonocardiales |  |
| S085 | Kryptoniales | Kryptoniales | Actinomarinales |  |
| Obscuribacteriales | Chthonomonadales | Chthonomonadales | Rickettsiales |  |
|  | SJA.28 | Thermoanaerobaculales | Paenibacillales |  |
|  | Paenibacillales | PLTA13 | Alicyclobacillales |  |
|  | Propionibacteriales | Bryobacteriales | Xanthomonadales |  |
|  | S085 | Rickettsiales | SJA.28 |  |
|  | PLTA13 | Elsterales | Chitinophagales |  |
|  | Isosphaerales |  | Ardenticatenales |  |
|  | Obscuribacteriales |  | Bacteroidales |  |
|  | Rokubacteriales |  | Gemmatimonadales |  |
|  | Solibacteriales |  | Nitrospirales |  |
|  |  |  | Rhizobiales |  |
|  |  |  | Defluviicoccales |  |
|  |  |  | CCD24 |  |
|  |  |  | Solirubrobacteriales |  |
|  |  |  | S085 |  |

**Table 14:**
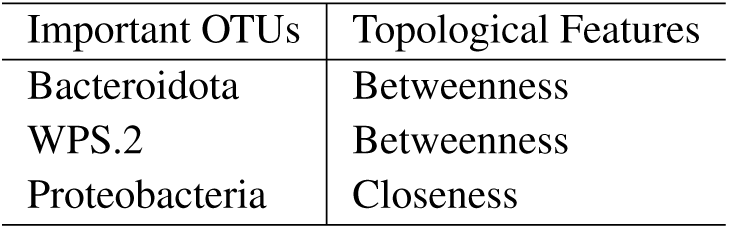
Important OTUs identified as key features in response to pitted scab at the **Phylum level using Strategy 1: Differential Centrality Analysis.** First, microbiome networks were constructed separately for ‘scab-infected tubers’ and ‘clean tubers’ using four inference methods: `SE_glasso`, `SPRING`, `SPARCC`, and `CMIMN`. Then, centrality metrics (Degree, Betweenness, Closeness, Eigenvector, and PageRank) were calculated for both networks, and the differences in centrality values between diseased and healthy conditions were computed for each method. OTUs ranked in the top 20% based on these centrality differences across all four methods were selected as important. The table presents OTUs that were consistently identified as important across all four inference methods. The Features column indicates the centrality measure that determined the significance of each OTU. These OTUs represent microbial taxa whose network connectivity consistently exhibited significant differences between healthy and diseased conditions across all four methods.

**Table 15:** Important OTUs identified as key features in response to pitted scab at the **Class level using Strategy 1: Differential Centrality Analysis.** First, microbiome networks were constructed separately for ‘scab-infected tubers’ and ‘clean tubers’ using four inference methods: `SE_glasso`, `SPRING`, `SPARCC`, and `CMIMN`. Then, centrality metrics (Degree, Betweenness, Closeness, Eigenvector, and PageRank) were calculated for both networks, and the differences in centrality values between diseased and healthy conditions were computed for each method. OTUs ranked in the top 20% based on these centrality differences across all four methods were selected as important. The table presents OTUs that were consistently identified as important across all four inference methods. The Features column indicates the centrality measure that determined the significance of each OTU. These OTUs represent microbial taxa whose network connectivity consistently exhibited significant differences between healthy and diseased conditions across all four methods.

| Important OTUs | Topological Features |
| --- | --- |
| Desulfotobacteriia | degree, Eigenvector Centrality, page_rank |
| Actinobacteria, AD3 | betweenness, closeness |
| Syntrophobacteria | Eigenvector Centrality |

**Table 16:** Important OTUs identified as key features in response to pitted scab at the **Order level using Strategy 1: Differential Centrality Analysis.** First, microbiome networks were constructed separately for ‘scab-infected tubers’ and ‘clean tubers’ using four inference methods: `SE_glasso`, `SPRING`, `SPARCC`, and `CMIMN`. Then, centrality metrics (Degree, Betweenness, Closeness, Eigenvector, and PageRank) were calculated for both networks, and the differences in centrality values between diseased and healthy conditions were computed for each method. OTUs ranked in the top 20% based on these centrality differences across all four methods were selected as important. The table presents OTUs that were consistently identified as important across all four inference methods. The Features column indicates the centrality measure that determined the significance of each OTU. These OTUs represent microbial taxa whose network connectivity consistently exhibited significant differences between healthy and diseased conditions across all four methods.

| Important OTUs | Topological Features |
| --- | --- |
| C0119 | Degree, Betweenness, Closeness, Eigenvector Centrality, Page Rank |
| Defluviicoccales | Closeness, Eigenvector Centrality |
| Bacteroidales | Closeness, Eigenvector Centrality |
| Kryptoniales | Eigenvector Centrality |
| B12.WMSP1 | Eigenvector Centrality |
| Desulfotobacteriales | Page Rank |

**Table 17:**
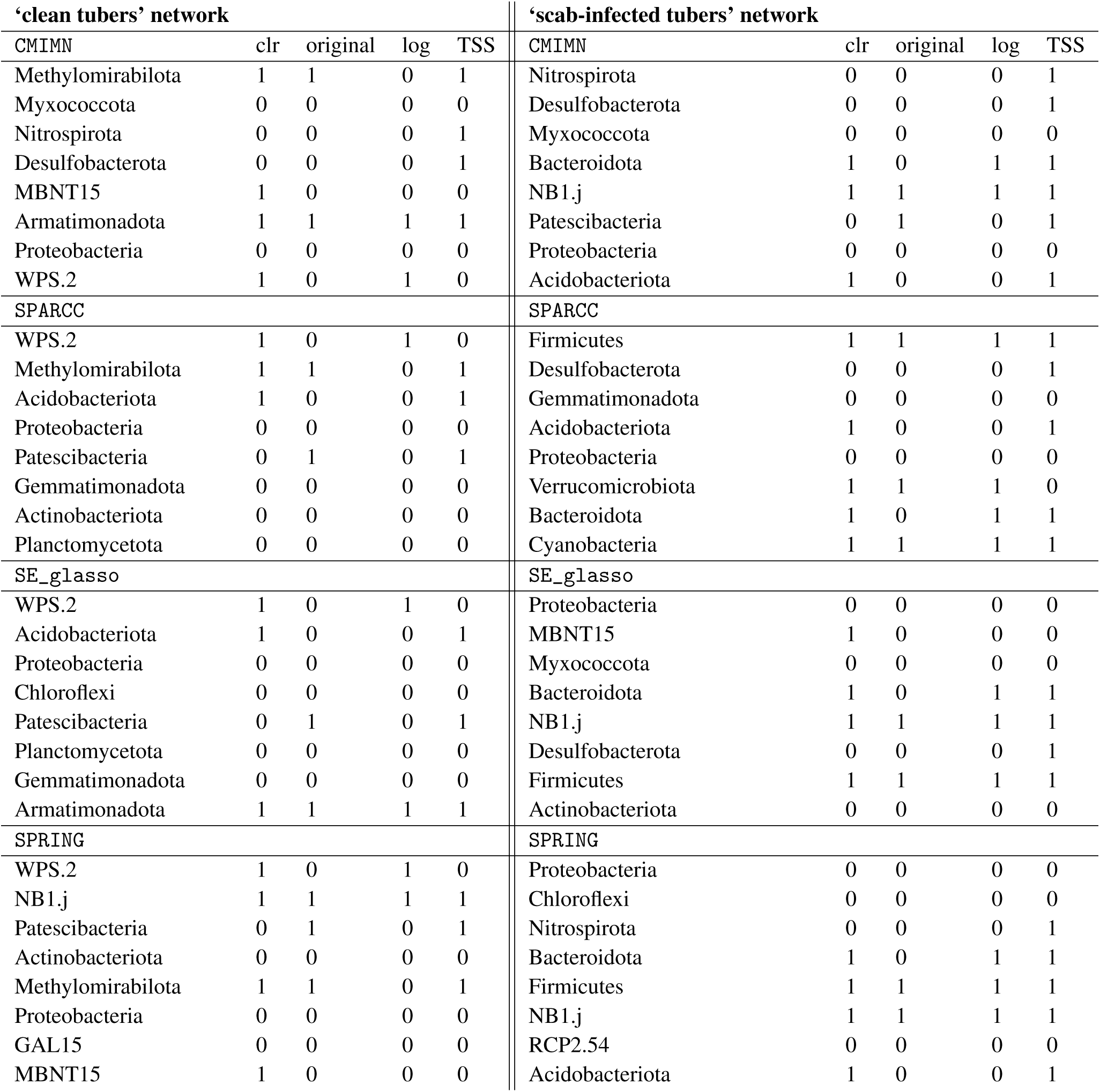
Selection of key Operational Taxonomic Units (OTUs) at the **Phylum** level in microbiome networks constructed separately for ‘clean tubers’ (left panel) and ‘scab-infected tubers’ (right panel) using network-based feature selection (Strategy 2: Composite Scoring Approach). **Steps for Identifying These OTUs:** 1-Network Construction: Microbiome networks were separately built for clean tubers and scab-infected tubers using four inference methods: `SE_glasso`, `SPRING`, `SPARCC`, and `CMIMN`. 2-Weighted Scoring Within Each Method: A weighted score was assigned to each OTU within each method based on multiple centrality metrics (Degree, Betweenness, Closeness, Eigenvector, and PageRank). Selection of Important OTUs: The top 20% of OTUs with the highest Score 1 were selected within each individual method. **Table Column Explanations:** First column in each panel: OTUs selected by Strategy 2 for each specific network inference method (i.e., these OTUs are among the top 20% highest-scoring OTUs for that method). Next four columns (CLR, Original, Log, TSS): Overlap between Strategy 2-selected OTUs and those identified by ML-based feature selection under different normalization approaches. A 1 in a column means that the ML method also identified the OTU as important under that normalization method. A 0 in a column means that the OTU was not selected by the ML method under that normalization method. Note: This table presents the weighted score for each OTU within each inference method. Unlike later steps in Strategy 2, this table does not include the combined score across all methods.

**Table 18:**
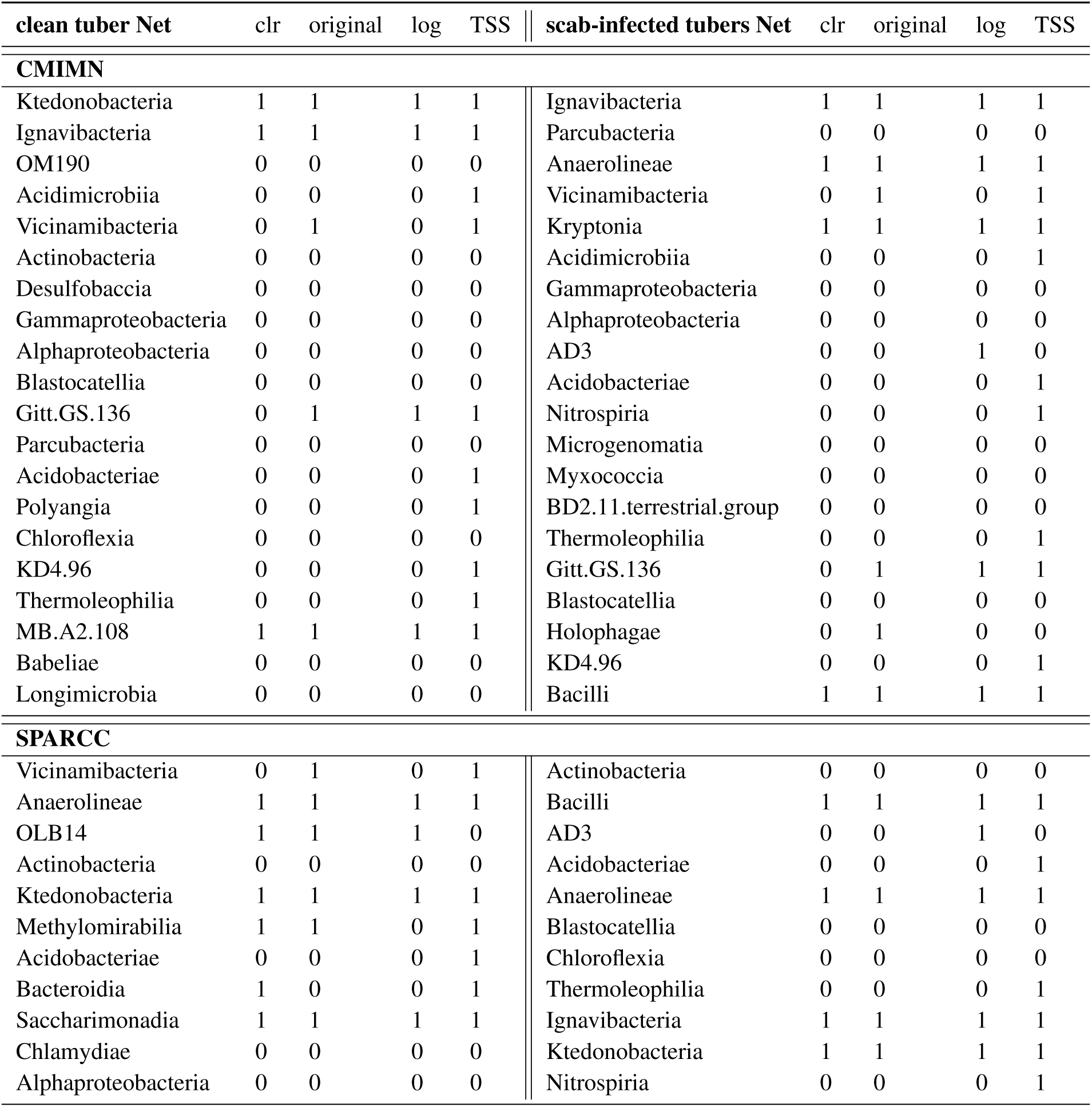
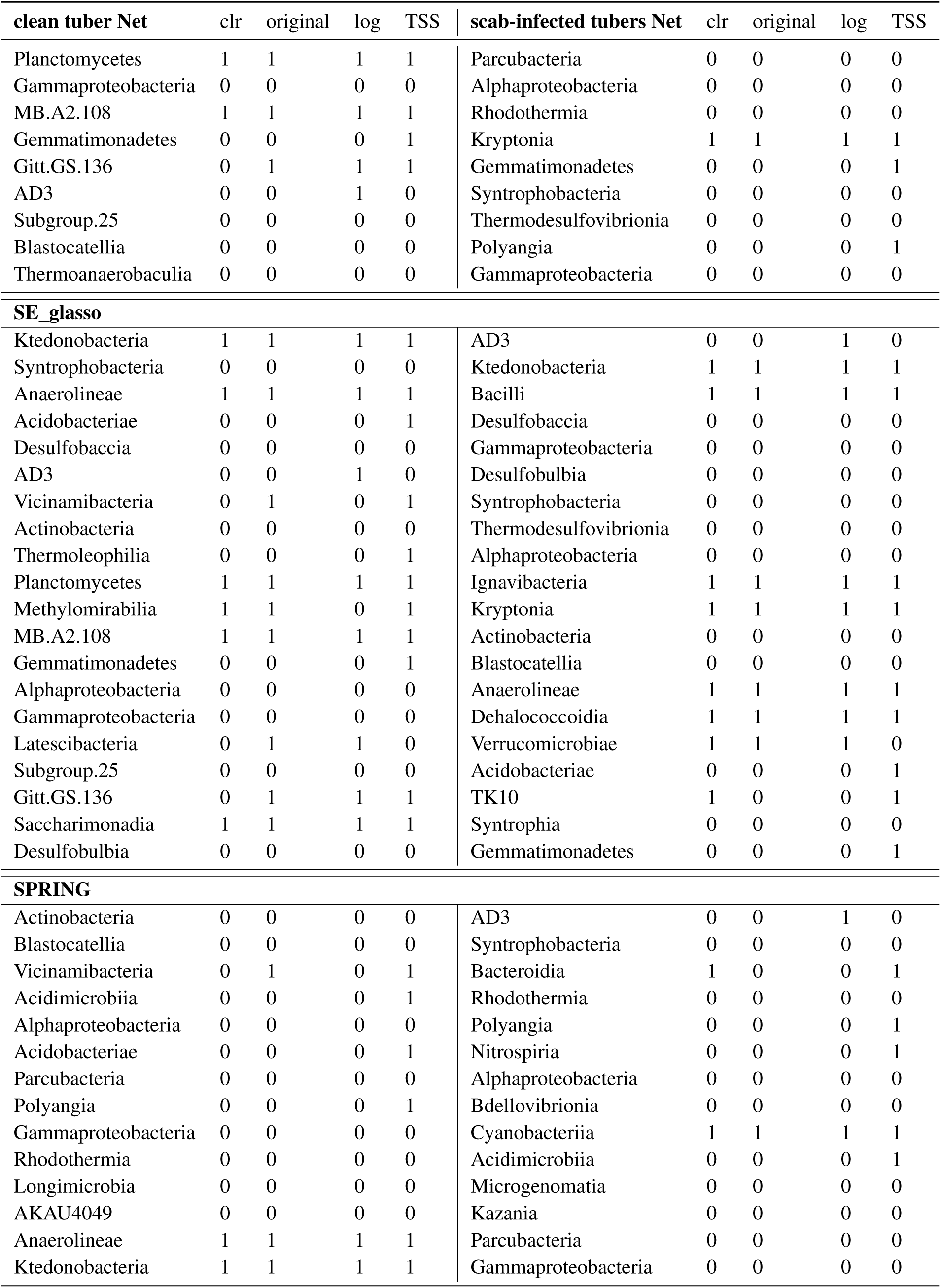
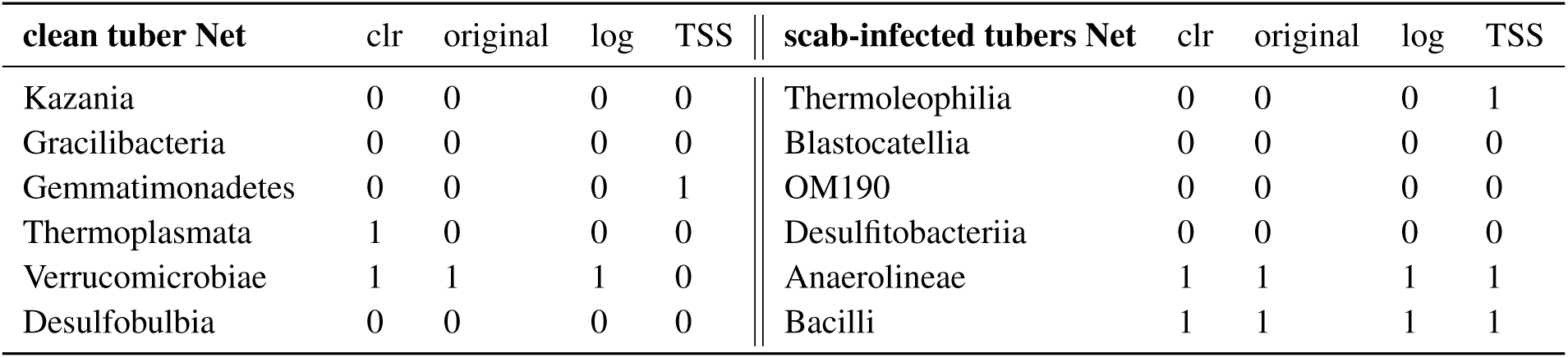
Selection of key Operational Taxonomic Units (OTUs) at the **Class** level in microbiome networks constructed separately for ‘clean tubers’ (left panel) and ‘scab-infected tubers’ (right panel) using network-based feature selection (Strategy 2: Composite Scoring Approach). **Steps for Identifying These OTUs:** 1-Network Construction: Microbiome networks were separately built for clean tubers and scab-infected tubers using four inference methods: `SE_glasso`, `SPRING`, `SPARCC`, and `CMIMN`. 2-Weighted Scoring Within Each Method: A weighted score was assigned to each OTU within each method based on multiple centrality metrics (Degree, Betweenness, Closeness, Eigenvector, and PageRank). Selection of Important OTUs: The top 20% of OTUs with the highest Score 1 were selected within each individual method. **Table Column Explanations:** First column in each panel: OTUs selected by Strategy 2 for each specific network inference method (i.e., these OTUs are among the top 20% highest-scoring OTUs for that method). Next four columns (CLR, Original, Log, TSS): Overlap between Strategy 2-selected OTUs and those identified by ML-based feature selection under different normalization approaches. A 1 in a column means that the ML method also identified the OTU as important under that normalization method. A 0 in a column means that the OTU was not selected by the ML method under that normalization method. Note: This table presents the weighted score for each OTU within each inference method. Unlike later steps in Strategy 2, this table does not include the combined score across all methods.

**Table 19:**
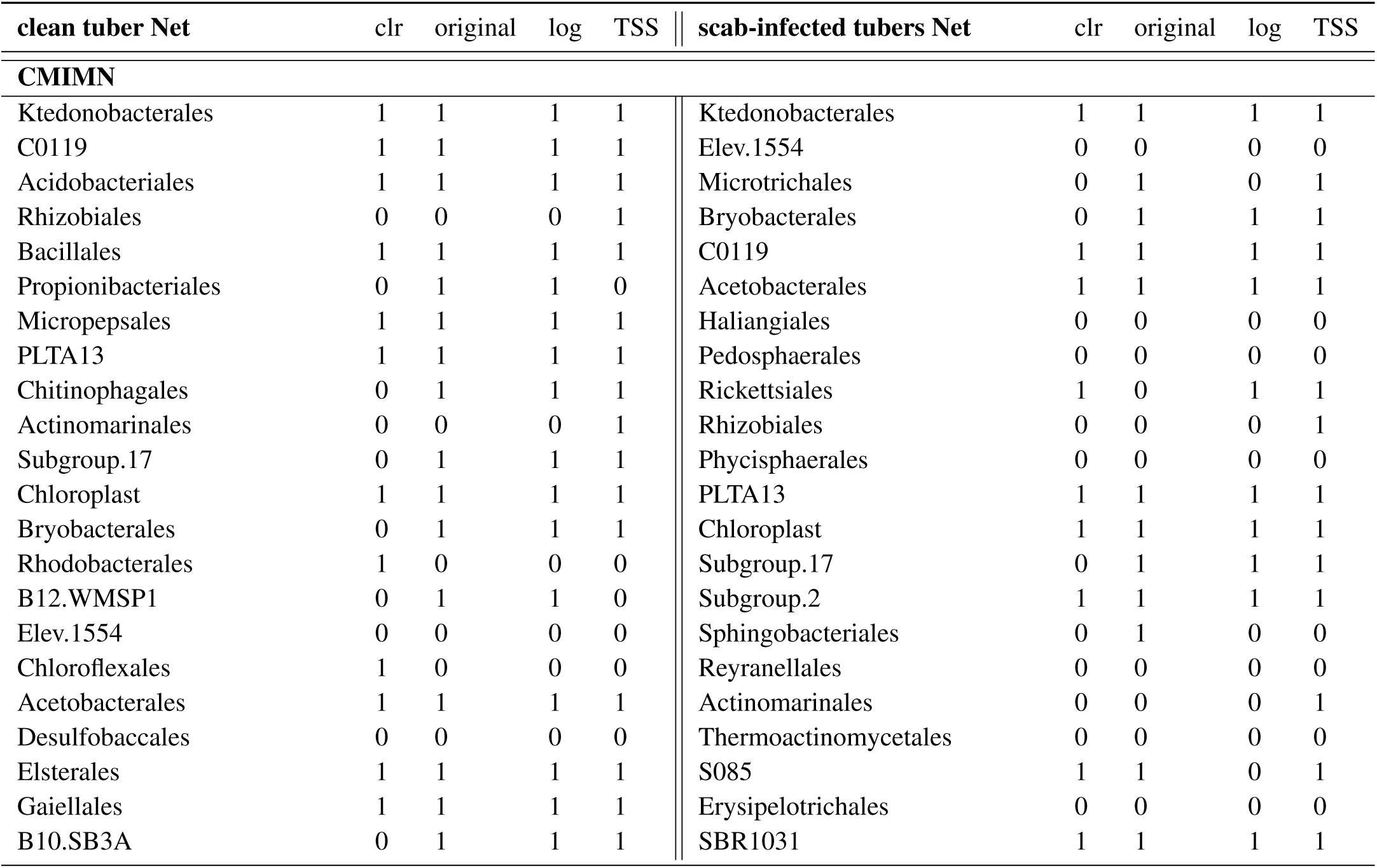
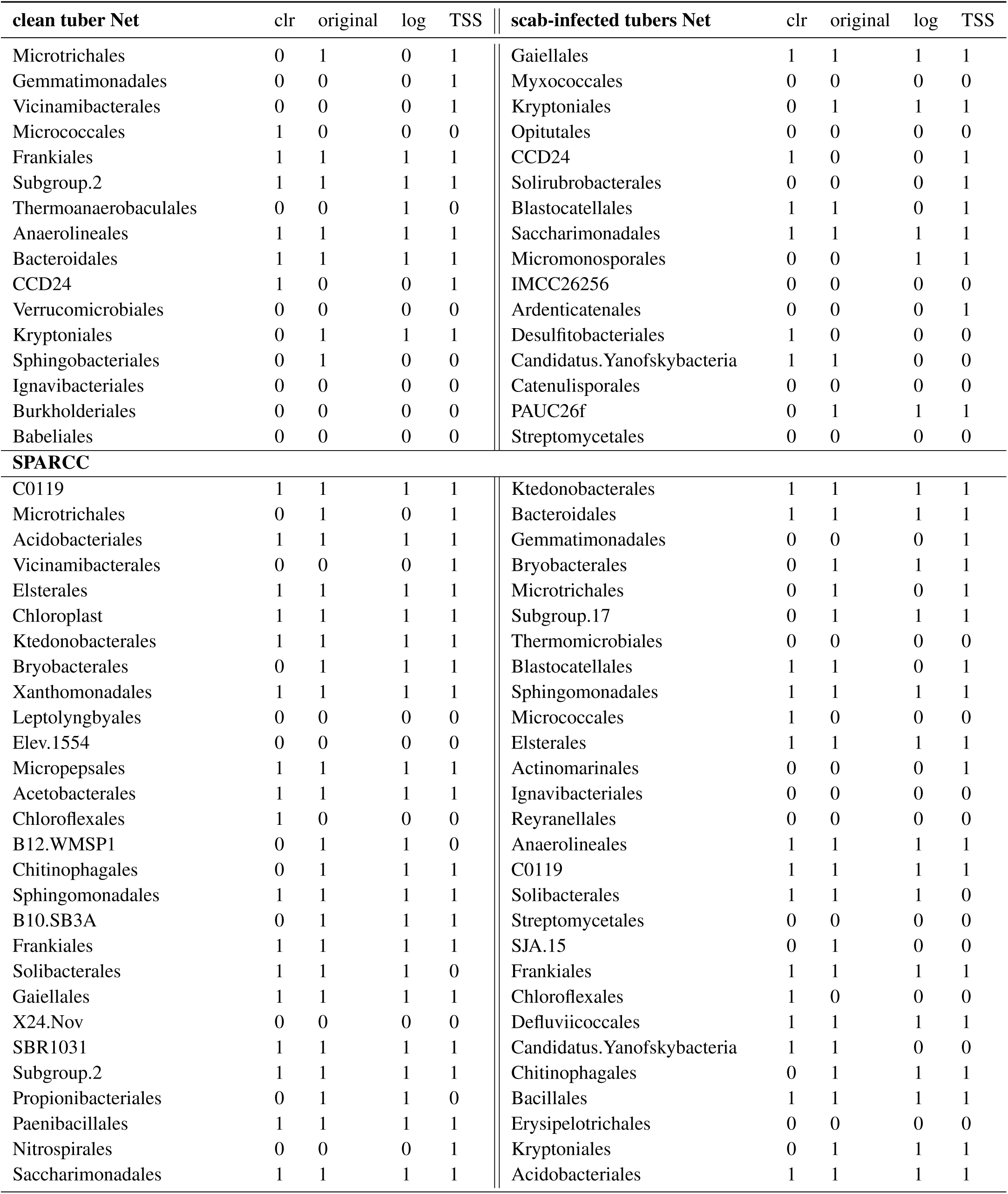
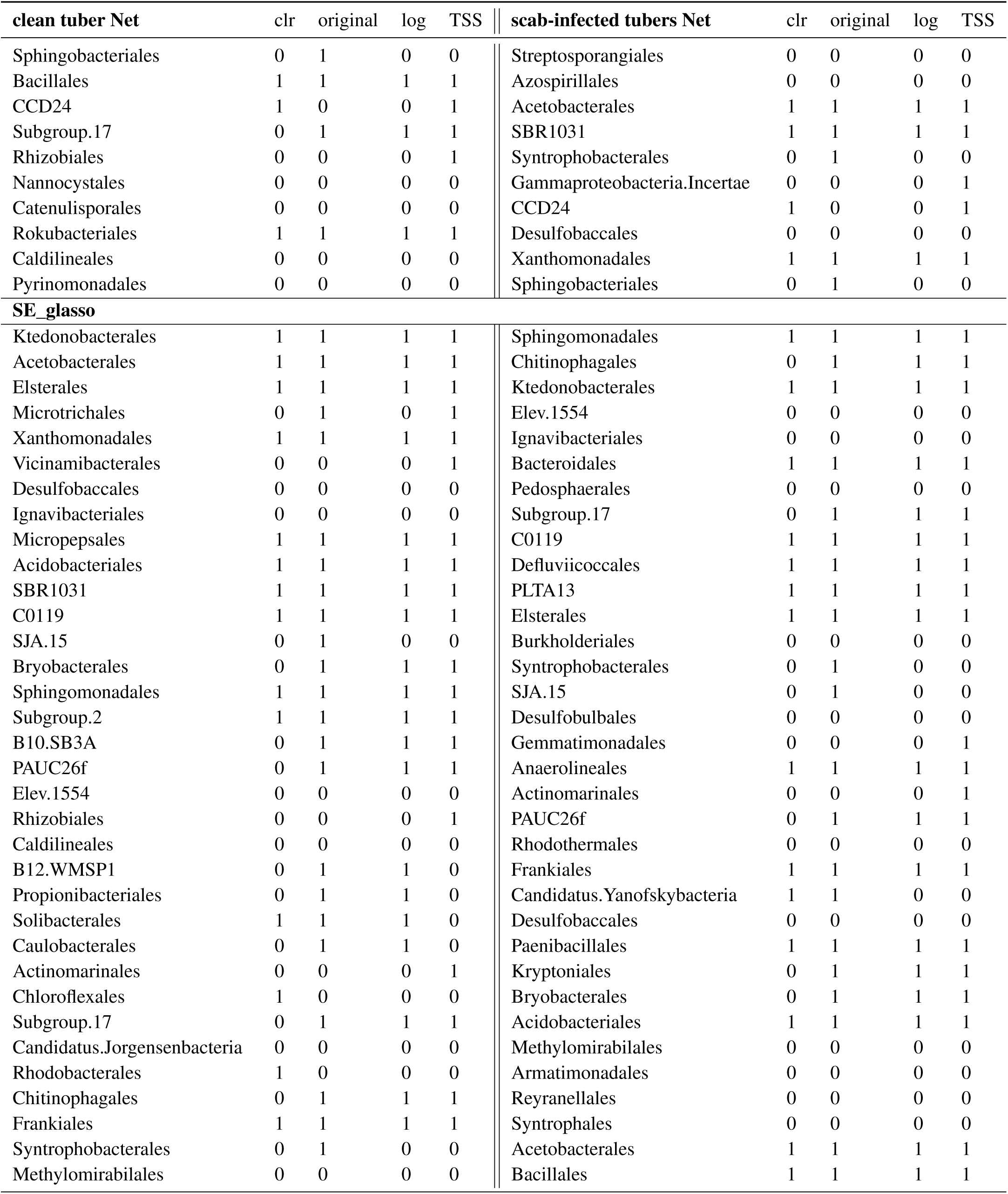
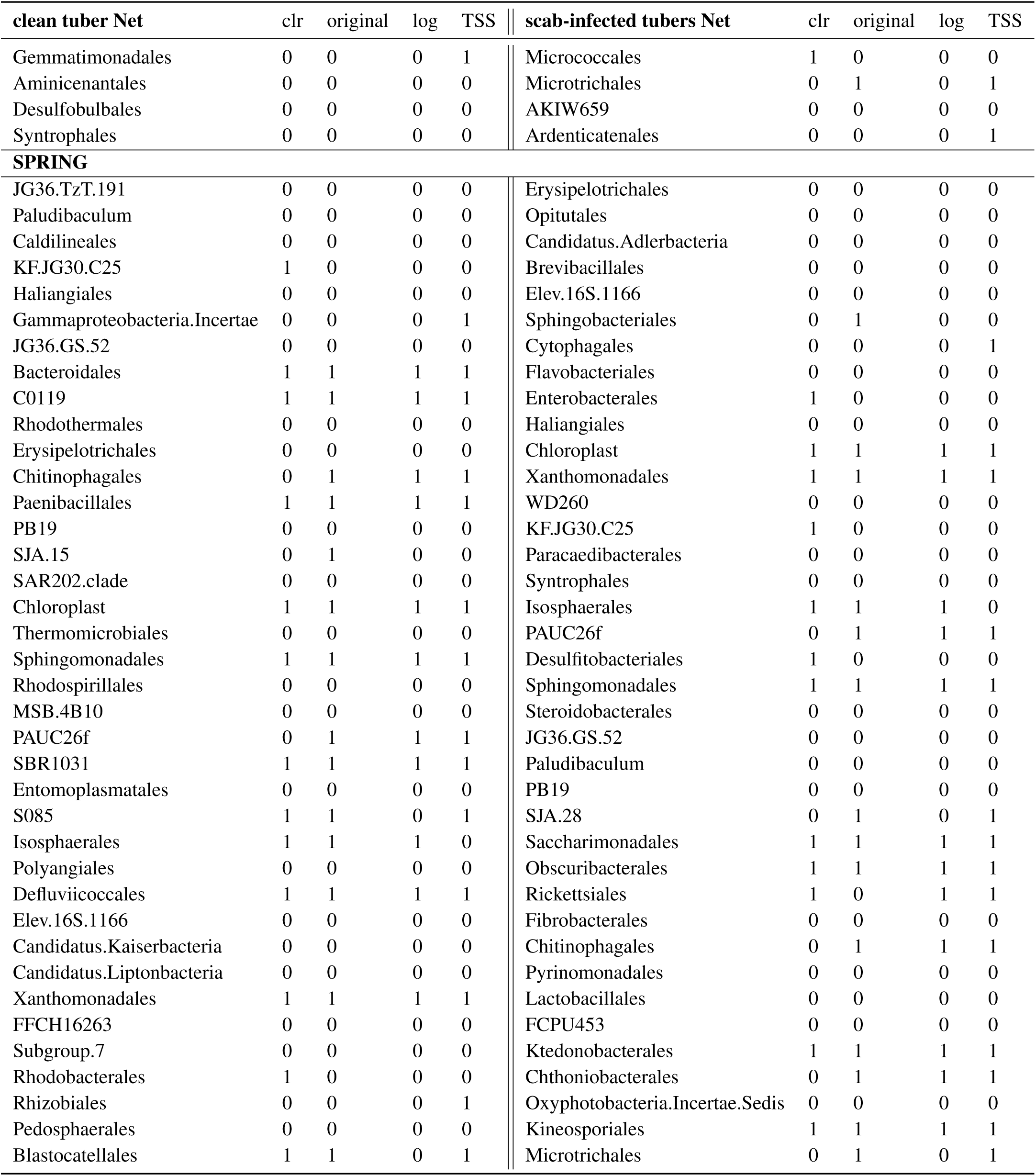
Selection of key Operational Taxonomic Units (OTUs) at the **Order** level in microbiome networks constructed separately for ‘clean tubers’ (left panel) and ‘scab-infected tubers’ (right panel) using network-based feature selection (Strategy 2: Composite Scoring Approach). **Steps for Identifying These OTUs:** 1-Network Construction: Microbiome networks were separately built for clean tubers and scab-infected tubers using four inference methods: `SE_glasso`, `SPRING`, `SPARCC`, and `CMIMN`. 2-Weighted Scoring Within Each Method: A weighted score was assigned to each OTU within each method based on multiple centrality metrics (Degree, Betweenness, Closeness, Eigenvector, and PageRank). Selection of Important OTUs: The top 20% of OTUs with the highest Score 1 were selected within each individual method. **Table Column Explanations:** First column in each panel: OTUs selected by Strategy 2 for each specific network inference method (i.e., these OTUs are among the top 20% highest-scoring OTUs for that method). Next four columns (CLR, Original, Log, TSS): Overlap between Strategy 2-selected OTUs and those identified by ML-based feature selection under different normalization approaches. A 1 in a column means that the ML method also identified the OTU as important under that normalization method. A 0 in a column means that the OTU was not selected by the ML method under that normalization method. Note: This table presents the weighted score for each OTU within each inference method. Unlike later steps in Strategy 2, this table does not include the combined score across all methods.

**Figure S4:**
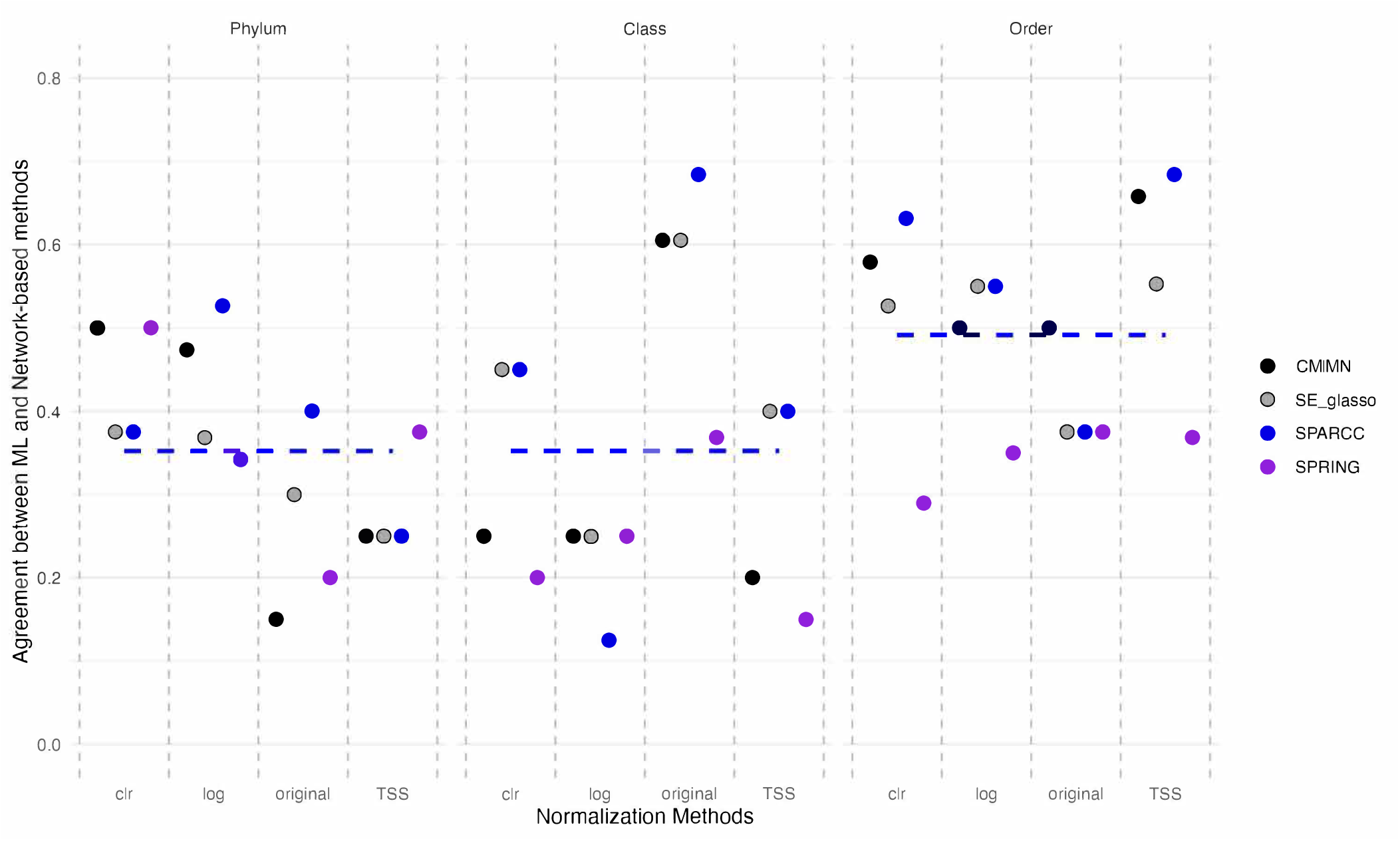
Average Overlap Between Machine Learning (ML) Methods based on different nomalized data sets (clr, original data, log transformation, and TSS transformation) and Network-Based Approaches for ‘clean tubers’ network in Phylum, Class, and Order levels.

**Figure S5:**
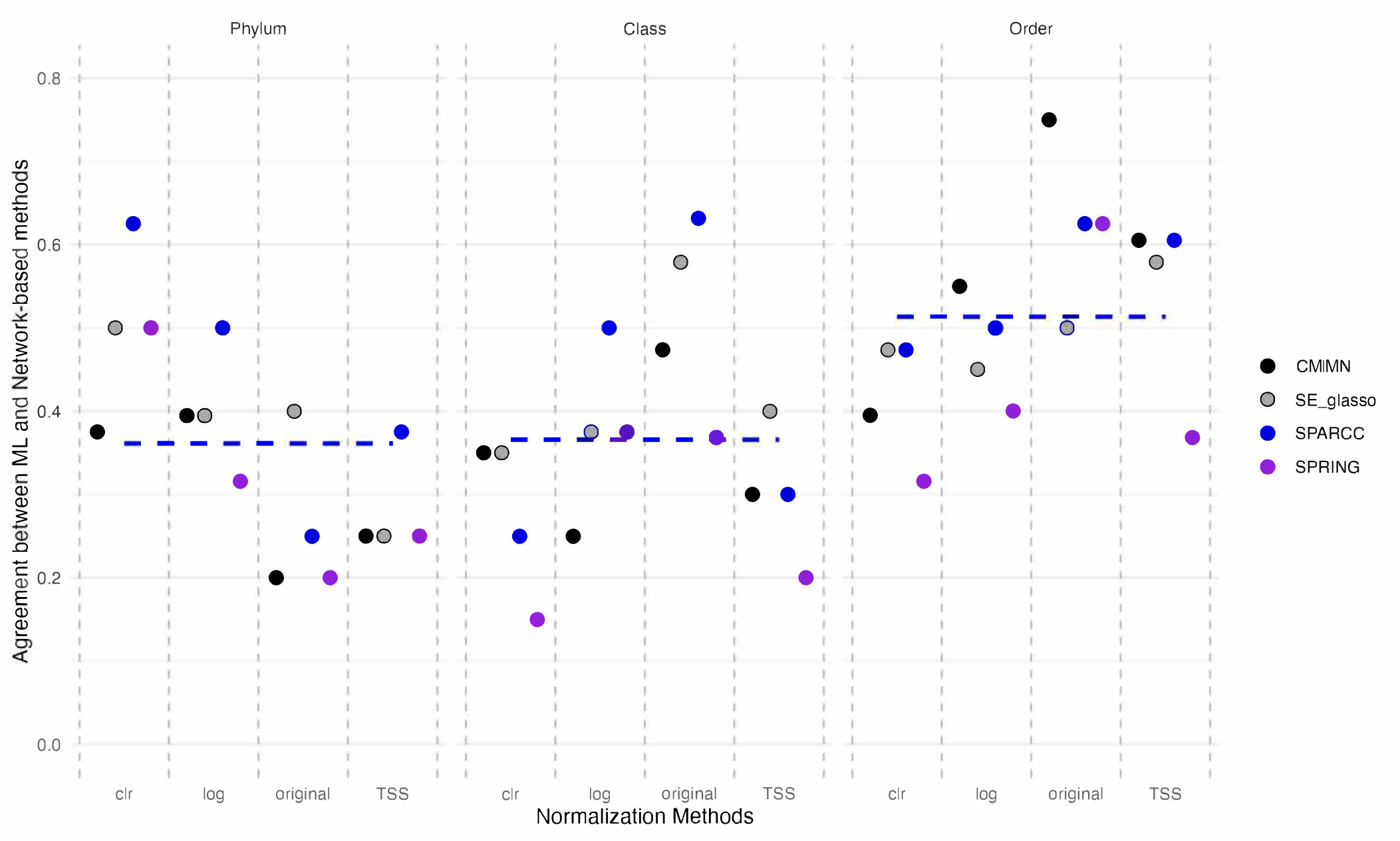
Average Overlap Between Machine Learning (ML) Methods based on different nomalized data sets (clr, original data, log transformation, and TSS transformation) and Network-Based Approaches for ‘scab-infected tubers’ network in Phylum, Class, and Order levels.

**Table 20:** Selection of Operational Taxonomic Units (OTUs) at the **Phylum** level in both networks of ‘Clean Tubers’ and ‘Scab-Infected Tubers’ using Network-Based Method (Strategy 2) and Machine Learning (ML) methods. The left part presents results from `CMIMN` and `SE_glasso`, while the right part displays results from `SPARCC` and `SPRING` algorithms. The first column represents OTUs selected by the network-based method (Strategy 2), and columns 2 to 4 show the overlap between OTUs selected by Strategy 2 and those chosen by ML methods using different data normalization approaches (clr, original, log, and TSS). A “1” indicates that the ML method selected the respective OTU, while “0” signifies that the ML method did not select the respective OTU.

| CMIMN | clr | original | log | TSS | SPARCC | clr | original | log | TSS |
| --- | --- | --- | --- | --- | --- | --- | --- | --- | --- |
| Myxococcota | 0 | 0 | 0 | 0 | Acidobacteriota | 1 | 0 | 0 | 1 |
| Nitrospirota | 0 | 0 | 0 | 1 | Proteobacteria | 0 | 0 | 0 | 0 |
| Desulfobacterota | 0 | 0 | 0 | 1 | Gemmatimonadota | 0 | 0 | 0 | 0 |
| Proteobacteria | 0 | 0 | 0 | 0 |  |  |  |  |  |
| SE_glasso | clr | original | log | TSS | SPRING | clr | original | log | TSS |
| Proteobacteria | 0 | 0 | 0 | 0 | NB1.j | 1 | 1 | 1 | 1 |

**Table 21:** Selection of Operational Taxonomic Units (OTUs) at the **Class** level in both networks of ‘Clean Tubers’ and ‘Scab-Infected Tubers’ using Network-Based Method (Strategy 2) and Machine Learning (ML) methods. The left part presents results from `CMIMN` and `SE_glasso`, while the right part displays results from `SPARCC` and `SPRING` algorithms. The first column represents OTUs selected by the network-based method (Strategy 2), and columns 2 to 4 show the overlap between OTUs selected by Strategy 2 and those chosen by ML methods using different data normalization approaches (clr, original, log, and TSS). A “1” indicates that the ML method selected the respective OTU, while “0” signifies that the ML method did not select the respective OTU.

| CMIMN | clr | original | log | TSS | SPARCC | clr | original | log | TSS |
| --- | --- | --- | --- | --- | --- | --- | --- | --- | --- |
| Ignavibacteria | 1 | 1 | 1 | 1 | Anaerolineae | 1 | 1 | 1 | 1 |
| Acidimicrobiia | 0 | 0 | 0 | 1 | Actinobacteria | 0 | 0 | 0 | 0 |
| Vicinamibacteria | 0 | 1 | 0 | 1 | Ktedonobacteria | 1 | 1 | 1 | 1 |
| Gammaproteobacteria | 0 | 0 | 0 | 0 | Acidobacteriae | 0 | 0 | 0 | 1 |
| Alphaproteobacteria | 0 | 0 | 0 | 0 | Alphaproteobacteria | 0 | 0 | 0 | 0 |
| Blastocatellia | 0 | 0 | 0 | 0 | Gammaproteobacteria | 0 | 0 | 0 | 0 |
| Gitt.GS.136 | 0 | 1 | 1 | 1 | Gemmatimonadetes | 0 | 0 | 0 | 1 |
| Parcubacteria | 0 | 0 | 0 | 0 | AD3 | 0 | 0 | 1 | 0 |
| Acidobacteriae | 0 | 0 | 0 | 1 | Blastocatellia | 0 | 0 | 0 | 0 |
| KD4.96 | 0 | 0 | 0 | 1 |  |  |  |  |  |
| Thermoleophilia | 0 | 0 | 0 | 1 |  |  |  |  |  |
| SE_glasso |  |  |  |  | SPRING |  |  |  |  |
| Ktedonobacteria | 1 | 1 | 1 | 1 | Blastocatellia | 0 | 0 | 0 | 0 |
| Syntrophobacteria | 0 | 0 | 0 | 0 | Acidimicrobiia | 0 | 0 | 0 | 1 |
| Anaerolineae | 1 | 1 | 1 | 1 | Alphaproteobacteria | 0 | 0 | 0 | 0 |
| Acidobacteriae | 0 | 0 | 0 | 1 | Parcubacteria | 0 | 0 | 0 | 0 |
| Desulfobaccia | 0 | 0 | 0 | 0 | Polyangia | 0 | 0 | 0 | 1 |
| AD3 | 0 | 0 | 1 | 0 | Gammaproteobacteria | 0 | 0 | 0 | 0 |
| Actinobacteria | 0 | 0 | 0 | 0 | Rhodothermia | 0 | 0 | 0 | 0 |
| Gemmatimonadetes | 0 | 0 | 0 | 1 | Anaerolineae | 1 | 1 | 1 | 1 |
| Alphaproteobacteria | 0 | 0 | 0 | 0 | Kazania | 0 | 0 | 0 | 0 |
| Gammaproteobacteria | 0 | 0 | 0 | 0 |  |  |  |  |  |
| Desulfobulbia | 0 | 0 | 0 | 0 |  |  |  |  |  |

**Table 22:** Selection of Operational Taxonomic Units (OTUs) at the **Order** level in both networks of ‘Clean Tubers’ and ‘Scab-Infected Tubers’ using Network-Based Method (Strategy 2) and Machine Learning (ML) methods. The left part presents results from `CMIMN` and `SE_glasso`, while the right part displays results from `SPARCC` and `SPRING` algorithms. The first column represents OTUs selected by the network-based method (Strategy 2), and columns 2 to 4 show the overlap between OTUs selected by Strategy 2 and those chosen by ML methods using different data normalization approaches (clr, original, log, and TSS). A “1” indicates that the ML method selected the respective OTU, while “0” signifies that the ML method did not select the respective OTU.

| CMIMN | clr | original | log | TSS | SPARCC | clr | original | log | TSS |
| --- | --- | --- | --- | --- | --- | --- | --- | --- | --- |
| Ktedonobacterales | 1 | 1 | 1 | 1 | C0119 | 1 | 1 | 1 | 1 |
| C0119 | 1 | 1 | 1 | 1 | Microtrichales | 0 | 1 | 0 | 1 |
| Rhizobiales | 0 | 0 | 0 | 1 | Acidobacteriales | 1 | 1 | 1 | 1 |
| PLTA13 | 1 | 1 | 1 | 1 | Elsterales | 1 | 1 | 1 | 1 |
| Actinomarinales | 0 | 0 | 0 | 1 | Ktedonobacterales | 1 | 1 | 1 | 1 |
| Subgroup.17 | 0 | 1 | 1 | 1 | Bryobacteriales | 0 | 1 | 1 | 1 |
| Chloroplast | 1 | 1 | 1 | 1 | Xanthomonadales | 1 | 1 | 1 | 1 |
| Bryobacteriales | 0 | 1 | 1 | 1 | Acetobacteriales | 1 | 1 | 1 | 1 |
| Elev.1554 | 0 | 0 | 0 | 0 | Chloroflexales | 1 | 0 | 0 | 0 |
| Acetobacteriales | 1 | 1 | 1 | 1 | Chitinophagales | 0 | 1 | 1 | 1 |
| Gaiellales | 1 | 1 | 1 | 1 | Sphingomonadales | 1 | 1 | 1 | 1 |
| Microtrichales | 0 | 1 | 0 | 1 | Frankiales | 1 | 1 | 1 | 1 |
| Subgroup.2 | 1 | 1 | 1 | 1 | Solibacteriales | 1 | 1 | 1 | 0 |
| CCD24 | 1 | 0 | 0 | 1 | SBR1031 | 1 | 1 | 1 | 1 |
| Kryptoniales | 0 | 1 | 1 | 1 | Sphingobacteriales | 0 | 1 | 0 | 0 |
| Sphingobacteriales | 0 | 1 | 0 | 0 | Bacillales | 1 | 1 | 1 | 1 |
|  |  |  |  |  | CCD24 | 1 | 0 | 0 | 1 |
|  |  |  |  |  | Subgroup.17 | 0 | 1 | 1 | 1 |
| SE_glasso | clr | original | log | TSS | SPRING | clr | original | log | TSS |
| Ktedonobacterales | 1 | 1 | 1 | 1 | Paludibaculum | 0 | 0 | 0 | 0 |
| Acetobacteriales | 1 | 1 | 1 | 1 | KF.JG30.C25 | 1 | 0 | 0 | 0 |
| Elsterales | 1 | 1 | 1 | 1 | Haliangiales | 0 | 0 | 0 | 0 |
| Microtrichales | 0 | 1 | 0 | 1 | JG36.GS.52 | 0 | 0 | 0 | 0 |
| Desulfobaccales | 0 | 0 | 0 | 0 | Erysipelotrichales | 0 | 0 | 0 | 0 |
| Ignavibacteriales | 0 | 0 | 0 | 0 | Chitinophagales | 0 | 1 | 1 | 1 |
| Acidobacteriales | 1 | 1 | 1 | 1 | PB19 | 0 | 0 | 0 | 0 |
| C0119 | 1 | 1 | 1 | 1 | Chloroplast | 1 | 1 | 1 | 1 |
| SJA.15 | 0 | 1 | 0 | 0 | Sphingomonadales | 1 | 1 | 1 | 1 |
| Bryobacteriales | 0 | 1 | 1 | 1 | PAUC26f | 0 | 1 | 1 | 1 |
| Sphingomonadales | 1 | 1 | 1 | 1 | Isosphaerales | 1 | 1 | 1 | 0 |
| PAUC26f | 0 | 1 | 1 | 1 | Elev.16S.1166 | 0 | 0 | 0 | 0 |
| Elev.1554 | 0 | 0 | 0 | 0 | Xanthomonadales | 1 | 1 | 1 | 1 |
| Actinomarinales | 0 | 0 | 0 | 1 |  |  |  |  |  |
| Subgroup.17 | 0 | 1 | 1 | 1 |  |  |  |  |  |
| Chitinophagales | 0 | 1 | 1 | 1 |  |  |  |  |  |
| Frankiales | 1 | 1 | 1 | 1 |  |  |  |  |  |
| Syntrophobacteriales | 0 | 1 | 0 | 0 |  |  |  |  |  |
| Methylomirabilales | 0 | 0 | 0 | 0 |  |  |  |  |  |
| Gemmatimonadales | 0 | 0 | 0 | 1 |  |  |  |  |  |
| Desulfobulbales | 0 | 0 | 0 | 0 |  |  |  |  |  |
| Syntrophales | 0 | 0 | 0 | 0 |  |  |  |  |  |

**Table 23:** Operational Taxonomic Units (OTUs) of significance in the ‘Clean Tubers’ network, selected by all four algorithms: CMIMN, SPARCC, SE_glasso, and SPRING.

| Level | important OTUs |  |  |  |  |  |
| --- | --- | --- | --- | --- | --- | --- |
| Phylum | Proteobacteria | WPS.2 |  |  |  |  |
| Class | Ktedonobacteria | Vicinamibacteria | Actinobacteria | Gammaproteobacteria | Alphaproteobacteria | Acidobacteriae |
| Order | C0119 | Rhizobiales | Chitinophagales |  |  |  |

**Table 24:**
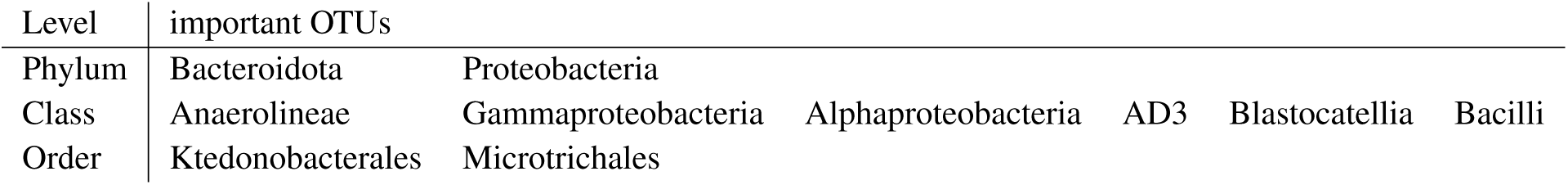
Operational Taxonomic Units (OTUs) of significance in the ‘scab-infected tubers’ network, selected by all four algorithms: CMIMN, SPARCC, SE_glasso, and SPRING.

| Level | important OTUs |  |  |  |  |  |
| --- | --- | --- | --- | --- | --- | --- |
| Phylum | Bacteroidota | Proteobacteria |  |  |  |  |
| Class | Anaerolineae | Gammaproteobacteria | Alphaproteobacteria | AD3 | Blastocatellia | Bacilli |
| Order | Ktedonobacterales | Microtrichales |  |  |  |  |

